# CNV-Finder: Streamlining Copy Number Variation Discovery

**DOI:** 10.1101/2024.11.22.624040

**Authors:** Nicole Kuznetsov, Kensuke Daida, Mary B. Makarious, Bashayer Al-Mubarak, Kajsa Atterling Brolin, Laksh Malik, Cedric Kouam, Breeana Baker, Raquel Real, Kathryn Step, Lara M. Lange, Lesley Wu, Miriam Ostrozovicova, Katherine M. Andersh, Pin-Jui Kung, Yasser Mecheri, Yi-Wen Tay, Behloul Soundous Malek, Nada Al Tassan, Maria Teresa Periñan, Samantha Hong, Mathew J. Koretsky, Lana Sargeant, Kristin Levine, Cornelis Blauwendraat, Kimberley J. Billingsley, Sara Bandres-Ciga, Hampton L. Leonard, Soraya Bardien, Huw R. Morris, Andrew B. Singleton, Mike A. Nalls, Dan Vitale, The Global Parkinson’s Genetics Program (GP2)

## Abstract

Copy Number Variations (CNVs) play pivotal roles in the etiology of complex diseases and are variable across diverse populations. Understanding the association between CNVs and disease susceptibility is significant in disease genetics research and often requires analysis of large sample sizes. One of the most cost-effective and scalable methods for detecting CNVs is based on normalized signal intensity values, such as Log R Ratio (LRR) and B Allele Frequency (BAF), from Illumina genotyping arrays. In this study, we present CNV-Finder, a novel pipeline integrating deep learning techniques on array data, specifically a Long Short-Term Memory (LSTM) network, to expedite the large-scale identification of CNVs within predefined genomic regions. This facilitates efficient prioritization of samples for time-consuming or costly subsequent analyses such as Multiplex Ligation-dependent Probe Amplification (MLPA), short-read, and long-read whole genome sequencing. We incorporate four genes to establish our methods—Parkin (*PRKN*), Leucine Rich Repeat And Ig Domain Containing 2 (*LINGO2*), Microtubule Associated Protein Tau (*MAPT*), and alpha-Synuclein (*SNCA*)—which may be relevant to neurological diseases such as Alzheimer’s disease (AD), Parkinson’s disease (PD), Progressive Supranuclear Palsy (PSP), or related disorders such as essential tremor (ET). By training our models on expert-annotated samples and validating them across diverse cohorts, including those from the Global Parkinson’s Genetics Program (GP2) and additional dementia-specific databases, we demonstrate the efficacy of CNV-Finder in accurately detecting deletions and duplications. Our pipeline outputs app-compatible files for visualization within CNV-Finder’s interactive web application. This interface enables researchers to review predictions and filter displayed samples by model prediction values, LRR range, and variant count in order to explore or confirm results. Our pipeline integrates this human feedback to enhance model performance and reduce false positive rates. Through a series of comprehensive analyses and validations using visual inspection, MLPA, short-read, and long-read sequencing data, we demonstrate the robustness and adaptability of CNV-Finder in identifying CNVs with regions of varied size, probe density, and noise. Our findings highlight the significance of contextual understanding and human expertise in enhancing the precision of CNV identification, particularly in complex genomic regions like 17q21.31. The CNV-Finder pipeline is a scalable, publicly available resource for the scientific community, available on GitHub (https://github.com/GP2code/CNV-Finder; DOI 10.5281/zenodo.14182563). CNV-Finder not only expedites accurate candidate identification but also significantly reduces the manual workload for researchers, enabling future targeted validation and downstream analyses in regions or phenotypes of interest.

## Introduction

### Copy Number Variation Detection in Disease Genetics

Copy Number Variations (CNVs) are structural variants (SVs) that play key roles in the diversity of genetic architecture and disease risk. Spanning hundreds to hundreds of thousands of base pairs, these genomic variations, including duplications, deletions, insertions, and complex rearrangements, deviate from the expected number of copies within DNA segments ^1^. Given the growing interest in understanding the contribution of SVs to disease heritability and susceptibility, these variants have been examined across multiple genetic modalities, including genotyping arrays, short-read, and long-read whole genome sequencing (WGS). Among these approaches, genotyping arrays are a cost-effective and widely accessible method for large-scale investigations of genetic data. CNV detection is facilitated by evaluating normalized signal intensity metrics, Log R Ratio (LRR) and B Allele Frequency (BAF) ^1, 2, 3^. The ability to detect CNVs and accurately determine their size depends on the genotyping array’s probe density across the region of interest. Enhancing the sensitivity and mitigating false positive rates of an array-based SV calling algorithm can expedite the identification of accurate candidates by significantly minimizing the number of samples that require laborious manual review. Here we present CNV-Finder, a tool that enables targeted screening with genotyping arrays, effectively pinpointing regions for future investigation via more specialized, expensive methods such as long-read sequencing.

### Genes of Interest

We selected suspected neurodegenerative disease (NDD) risk loci for model development, including Parkin (*PRKN;* chr6:161347557-162727802 [hg38]), Leucine Rich Repeat And Ig Domain Containing 2 (*LINGO2*; chr9:27948085-29213000 [hg38]), and Microtubule Associated Protein Tau (*MAPT*; chr17:45894381-46028333 [hg38]). These intervals exhibited sufficiently high rates of deletions and duplications to allow for the creation of large, robust training sets for both CNV types. The variants in these regions were well-defined, minimizing noise and ensuring consistency among manual classifications. Alpha-Synuclein (*SNCA*; chr4:89724098-89838296 [hg38]) was included to demonstrate the final models’ usability on previously unseen genes with markedly lower frequencies.

Bi-allelic *PRKN* structural variants can cause autosomal recessive Parkinson’s disease (PD). However, the significance of common single heterozygous duplications and deletions in PD risk remains uncertain due to incomplete sequence and CNV data across case and control populations, as well as conflicting findings in literature ^4,5,6^. Previous literature has also indicated potential disease-association between *LINGO2* SVs and conditions such as essential tremor (ET) and PD, with similar limitations due to sample size ^7, 8^. Additionally, *MAPT* represents a particularly complex genomic region characterized by high linkage disequilibrium, distinct functional haplotypes, and a common inversion observed among Europeans ^9^. This gene, along with neighboring genes such as *KANSL1* in chromosomal band 17q21.31, has attracted interest for potential associations with diseases including Progressive Supranuclear Palsy (PSP), Frontotemporal Dementia (FTD), Dementia with Lewy Bodies (DLB), and Alzheimer’s disease (AD) ^10, 11, 12^. Previous literature has supported an association between the H1-haplotype of *MAPT* with additional diseases such as PD and other tauopathies like Corticobasal Degeneration (CBD), which may present clinically as Corticobasal Syndrome (CBS) ^13, 14^. The *MAPT* H2 haplotype, on the other hand, has been associated with reduced risk in AD and explored for potential associations with other diseases ^15, 16, 17^. Triplications and duplications in *SNCA* have strong evidence primarily linking the gene to monogenic PD ^18, 19, 20^. Utilizing the scalability of our pipeline, we developed a publicly available resource for the scientific community to conduct large-scale, parallelized searches across multiple cohorts to further investigate the role of SVs in the genetic architecture of NDDs, across wide ranges of phenotypes and ancestries.

## Materials and Methods

### Genotyping Data Inputs

All samples featured in both model training and testing underwent genotyping using the Illumina NeuroBooster array (NBA) ^21^ and subsequent processing by the Global Parkinson’s Genetics Program (GP2) ^22^. The NBA is specifically optimized for NDDs, therefore improving coverage in the regions we are analyzing and enhancing reliability in signal intensity measures. Notably, around 1.9 million variants are included in the array, with over 95,000 variants specific to the context of neurological disorders. One file containing the position, chromosome, BAF, and LRR values for all single nucleotide-polymorphism (SNPs) serves as the necessary input for the pipeline. We incorporated all available probe intensity data, including instances where a single SNP was captured by multiple probes, as well as any CNV-specific probes on the array. Because these duplicate and CNV-targeted probes only make up a small fraction of the input file and our algorithm relies on aggregated measures across many variants, retaining only one probe per chromosome position has minimal impact on results. To enhance data quality, users have the option to specify a customizable GenTrain (https://www.illumina.com/content/dam/illumina/gcs/assembled-assets/marketing-literature/gentrain-tech-note-m-gl-01258/gentrain-tech-note-m-gl-01258.pdf) ^23^ score cut-off, which defaults to 0.2, thus filtering out SNPs with suboptimal calling quality. To observe the full range of potential predictions, no additional quality control was applied to the samples in our analysis; however, users may additionally upload a PLINK v1.9 (RRID:SCR_001757) ^24^ bim format file or PLINK v2.3 ^25^ pvar format file to limit samples and variants to those that passed quality control detailed elsewhere ^26^. Furthermore, users are required to specify either a gene name or chromosome, along with start and stop positions in base pairs, delineating the region of interest. All graphical representations and positional references in this paper adhere to the Genome Reference Consortium Human Build 38 (hg38) ^27^.

### Samples

All GP2 cohorts involved in model creation and evaluation are delineated in Supplementary Table 1 and can be searched on GP2’s Cohort Dashboard by their study abbreviations (https://gp2.org/cohort-dashboard-advanced/). Supplementary Table 2 presents the manually curated set of 184 samples with 156 PD cases and 28 controls that were used in training preliminary models for each CNV type. This array data underwent visual assessment by a team of 13 NDD research scientists to identify samples exhibiting deletions or duplications of any size within *PRKN*. The preliminary training set comprises 70 deletions (Supplementary Table 2a), 29 duplications (Supplementary Table 2b), and 85 samples lacking visible CNVs in *PRKN* (Supplementary Table 2c). Short-read WGS was used to confirm the absence of visible structural variants in negative samples. The Accelerating Medicines Partnership program for Parkinson’s disease (AMP-PD) short-read WGS data collection has been previously outlined by Iwaki et al ^28^ with additional details on SNV calling by Billingsley et al ^29^.

Our initial training set was iteratively expanded on by using CNV-Finder’s predictions for cohorts listed in Supplementary Table 3. This group of samples encompasses a broad spectrum of phenotypes and spans 11 ancestries to diversify our training sets. Figure 1 outlines the process by which samples were incorporated into preliminary, updated, and final models for each CNV type, following expert review of model predictions. Expert-annotations on samples from the Coriell Institute for Medical Research (CORIELL), detailed in Supplementary Table 4, were withheld during preliminary and updated model training to assess performance improvements between these two models. CORIELL samples were then incorporated into our largest, final training set.

**Figure 1.**
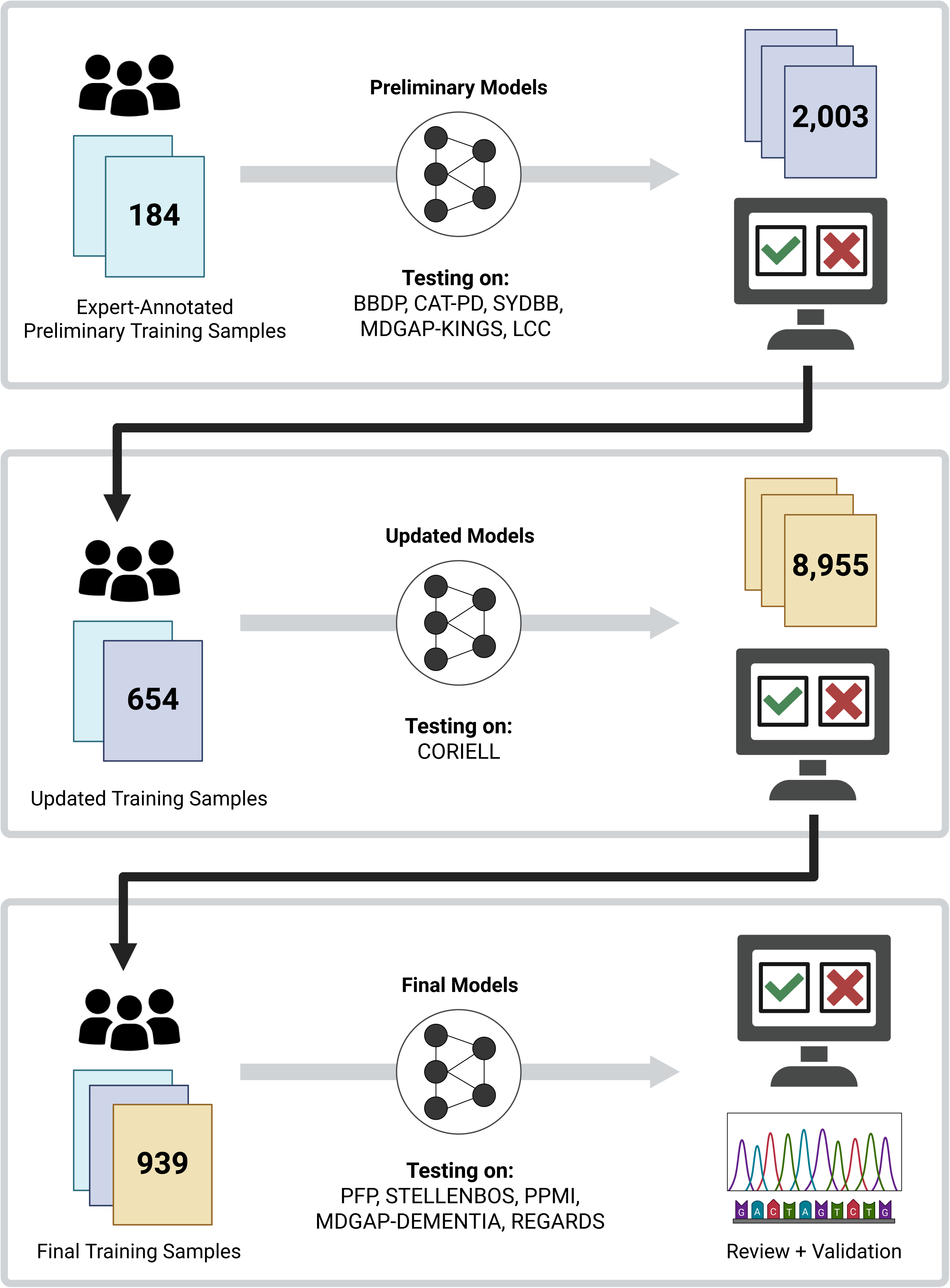
Workflow to demonstrate the iterative growth of our training sets through testing and prediction reviews on additional cohorts.

Multi-modal and extended datasets from additional studies provided further validation and insight into the performance of our final models. Samples from the Parkinson’s Families Project (PFP) ^30^ and Parkinson’s Disease in Southern Africa (STELLENBOS) ^31, 32, 33^, described in Supplementary Table 5, were used for Multiplex Ligation-dependent Probe Amplification (MLPA) validation of *PRKN* and *SNCA.* Parkinson’s Progression Markers Initiative (PPMI) samples highlighted in Supplementary Table 6 enabled short and long-read WGS validation of the three genes involved in model development. Finally, a subset of samples from the (MDGAP-DEMENTIA) and REasons for Geographic and Racial Differences in Stroke (REGARDS) studies in Supplementary Table 7 supported an in-depth exploration of the 17q21.31 region encompassing *MAPT*, particularly within dementia-related phenotypes.

### Long-read Whole Genome Sequencing

DNA was extracted from frozen blood samples as detailed in the protocol available at dx.doi.org/10.17504/protocols.io.x54v9py8qg3e/v1. In summary, high molecular weight (HMW) DNA was isolated from 1mL of frozen whole blood using the KingFisher Apex instrument, following an adapted version of the PacBio Nanobind HT HMW DNA Extraction 1mL Whole Blood KingFisher Apex protocol, along with the Nanobind HT 1mL Blood Kit (102-762-800) from PacBio. The DNA underwent a size selection step with the PacBio Short Read Eliminator Kit (102-208-300) to eliminate fragments up to 25 kilobases (kb). Subsequently, the DNA was sheared to a target size of 30kb using the Megaruptor 3 instrument and the Diagenode Megaruptor 3 Shearing Kit (E07010003). The DNA library was then prepared using the SQK-LSK114 Ligation Sequencing Kit from Oxford Nanopore Technologies (ONT). The samples were loaded onto a PromethION R10 flow cell (FLO-PRO114M) according to ONT’s standard operating procedures and run for 72 hours on a PromethION device.

Fast5 files containing raw signal data were acquired from sequencing conducted with Oxford Nanopore Technologies’ MinKNOW v22.10.7 (https://nanoporetech.com/about-us/news/introducing-new-minknow-app). Super accuracy base calling was performed on all Fast5 files for each sample using Oxford Nanopore Technologies’ Dorado v0.6.0 (RRID:SCR_025883). Fastq files that passed quality control filters during the super accuracy basecalling step were subsequently aligned to the hg38 reference genome using Winnowmap (RRID:SCR_025349). The resulting SAM files were sorted, converted to BAM files, and indexed with SAMtools v1.20 (RRID:SCR_002105) ^34^. These BAM files were then merged, sorted, and indexed to generate a final BAM file for each sample. SVs were detected and genotyped using Sniffles2 v.2.3 (RRID:SCR_017619, https://github.com/fritzsedlazeck/Sniffles) ^35^.

#### Pipeline

##### Model Framework

Our pipeline employs a recurrent neural network known as Long Short-term Memory (LSTM) ^36^, known for its proficiency in capturing long-term dependencies within sequential data. We stack three LSTM layers, each passing its sequential representations to the next, forming a deeper network that learns multiple levels of abstraction for improved feature extraction. A hard sigmoid activation function was chosen for its decreased computational cost and success with deep learning binary classification tasks. Therefore, our model outputs predicted values between 0 and 1 which correspond to the likelihood of a CNV for that sample in the region of interest (Supplementary Figure 1).

We apply overlapping sliding windows to the signal intensity files during data preparation for our model. To determine window sizes for each gene or interval of interest, we utilize customizable variables for “split” and “total window counts” within the region. For instance, if a model is trained with a split of 5 and a total window count of 50, this signifies that a gene like *PRKN* is divided into 5 equal segments, each spanning 276,049 base pairs without any overlap. Subsequently, 50 windows are computed to overlap these segments by 22,534 bases as shown in Supplementary Figure 2. Similarly, in *MAPT*, the 5 equal intervals are 26,790 base pairs long with windows overlapping by 2,168 bases. This approach creates proportional windows to span any gene regardless of its size, without the need for padding with artificial bases to create sequences of equal lengths. Furthermore, it enables the application of a single model to all regions prepared with the same split and window counts. We also include a modifiable buffer that can be added to both sides of a region of interest before window calculation. In our analysis, we incorporated a default 250 kb buffer to facilitate the exploration of variant behavior surrounding the regions of interest. We later adjusted this metric to 10 megabases (Mb) for the analysis of *SNCA*, to better capture the scale of large SVs found in this risk locus.

Features summarized in Supplementary Table 8 are aggregated within and across each window to capture trends such as visible positional shifts in LRR or BAF and probabilistic dosage spikes. These spikes are depicted in Figure 2 by LRR or BAF against chromosome position (in base pairs) for *PRKN* deletions (Column A) and duplications (Column B). SNP variants in this figure, color-coded for visibility, are deemed “CNV candidates” if they fall within specific ranges recommended by the array manufacturer:

- Deletions: LRR < −0.2
- Duplications: LRR > 0.2
- Insertions: 0.15 < BAF < 0.35 or 0.65 < BAF < 0.85

**Figure 2.**
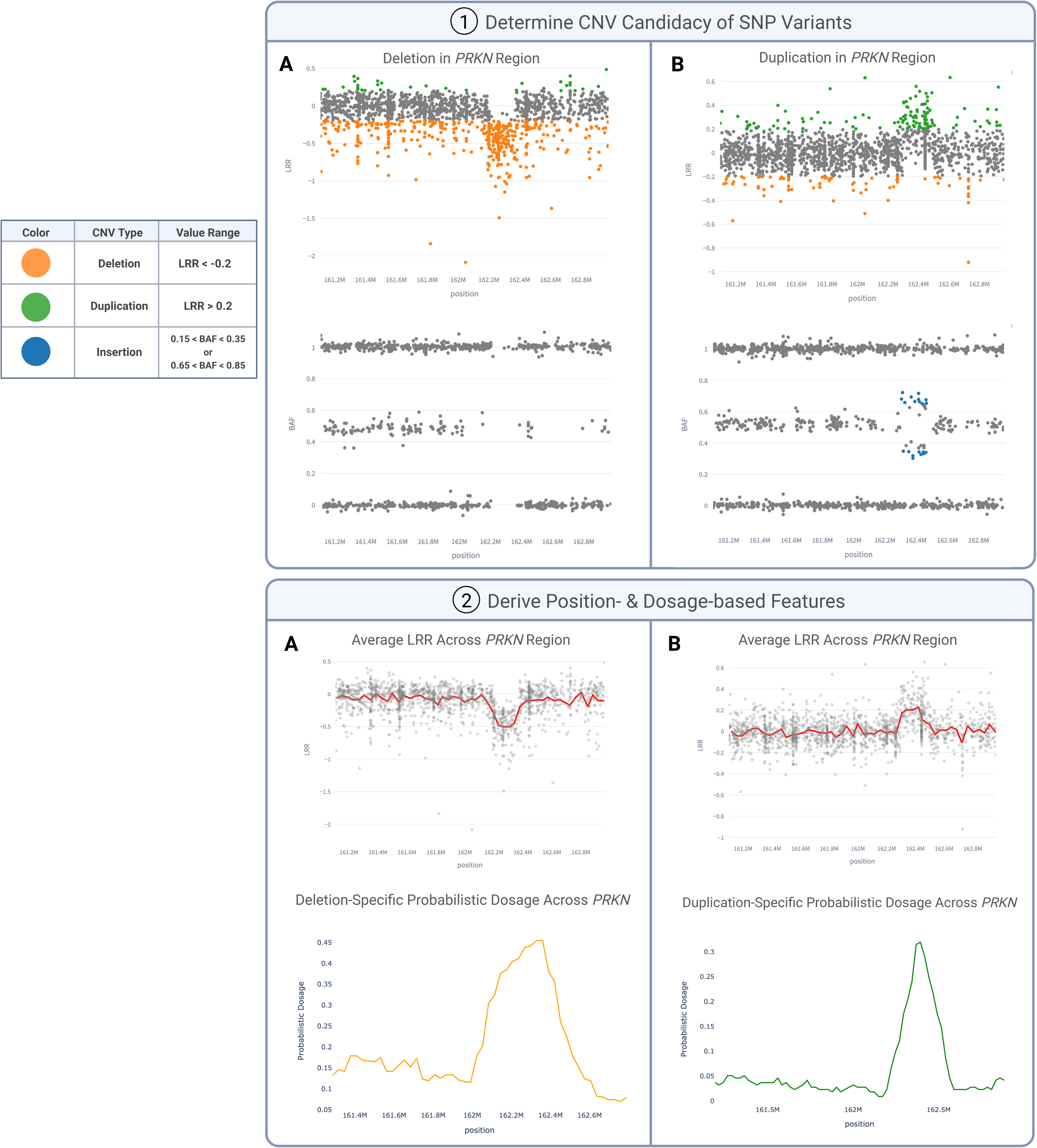
Feature creation to capture deletions (A) and duplications (B). (1) Illumina-based thresholds are applied to SNPs to determine if their LRR and BAF values fall into the proper ranges to be considered deletions and duplications. (2) Features in Supplementary Table 8 are calculated on the variants within these specified CNV ranges (CNV candidates).

To comprehensively capture the mid-range of BAFs, we incorporate additional features encompassing BAF values from 0.15 to 0.85, inclusive. This broader range of frequencies is crucial for capturing characteristics such as the diverging branching pattern consistently observed in duplications of varying size and BAF values ^37^. Most model features are only derived from the CNV candidates, aiming to enhance precision and reduce computational cost by subsetting the extensive pool of SNPs within each region.

A five-fold cross-validation was performed on the training set to ascertain optimal window sizes and feature combinations, listed in Supplementary Table 8, for predicting each CNV type. A split of 5, sliding window of 50, and the 4th feature combination, earned an average Area Under the Curve (AUC) (Supplementary Table 9) of 0.93 for our deletion model. For the duplication model, an average AUC of 0.92 was obtained with a split of 10, sliding window of 70, and the 6th feature combination. Therefore, if you choose to utilize our existing pre-trained models for a cohort of size N samples, the pipeline will reshape the model-ready input files to the necessary dimensions of [N, 50, 13] and [N, 70, 11] for the deletion and duplication models, respectively.

##### Semi-Automated Integration with Visualization

The final step of CNV-Finder prepares app-compatible files for visualization within our interactive web application. This interface enables researchers to review predictions and filter displayed samples by model prediction values, LRR range, and variant count in order to explore or confirm results. Users have the flexibility to examine results by selecting from options such as “Yes,” “Maybe,” or “No,” which are used to record chosen sample IDs, their respective regions of interest, and the type of CNV under evaluation. Logs generated by the application can subsequently serve as input to the pipeline for enlarging the training set or training a new model.

## Results

CNV-Finder facilitates machine learning preparation of genotyping data, supports visual assessment of predicted CNV-carriers through a web-application, and refines its models by incrementally building on training sets from initial expert annotations. For each CNV type, three models–preliminary, updated, and final–were trained using identical architectures, differing only in the size of their training sets. The preliminary model was applied to five cohorts (BBDP, CAT-PD, LCC, MDGAP-KINGS, SYDBB in Table 1) and samples that received predicted values equal to or exceeding 0.8 were reviewed. Based on this evaluation, 116 samples (31 deletions and 85 negative examples) were incorporated into the deletion training set, while 354 samples (132 duplications and 222 negative examples) were added to the duplication training set. At this stage, the samples with visible CNVs in the updated training sets were composed of 82% *PRKN* deletions, 18% *LINGO2* deletions, 79% duplications near *MAPT*, and 21% *PRKN* duplications.

The updated model was then applied to the large-scale CORIELL cohort, adding 159 samples (150 deletions, 9 negative samples) to the final deletion training set and 126 samples (53 duplications, 73 negative samples) to the final duplication training set. This resulted in final training sets composed of 71% *PRKN* deletions, 29% *LINGO2* deletions, 59% duplications near *MAPT*, and 41% *PRKN* duplications. Evaluation of the final model was conducted on PFP, STELLENBOS, and PPMI cohorts, with validation from MLPA, short and long-read whole genome sequencing.

Development of the preliminary, updated, and final models involved over 11,000 samples. Benchmarking was conducted on 720 samples within the BBDP cohort in the *PRKN* gene locus. All cohorts and genes can be processed in parallel, potentially benefiting from enhanced efficiency through multi-core computing or chromosome-specific input organization. Without requiring any pre-processing or input subsetting, the most computationally demanding steps—data processing and feature aggregation—produced a model-ready 16 MB file for 70 overlapping windows in approximately 23 minutes with 1 CPU and 7.5 minutes with 8 CPUs. Using 1 CPU, model testing on BBDP samples completed in 19 seconds, while training the final and largest duplication model on 594 samples generated a 1 MB model file in 31 seconds.

## Discussion

### Performance Improvements with Growing Training Sets

#### Visual Validation to Assess Iterative Training Sets

Even with its smallest training set, CNV-Finder correctly identified 97% of the expert-annotated CORIELL deletions and 84% of the duplications in *PRKN* with predicted values above 0.5 (Supplementary Table 10). Subsequent sample additions to the training set reduced false positives of the deletion model and decreased false negatives of the duplication model, while maintaining high rates of true positives in the updated model. Consequently, this improvement led to increases in the AUC metrics, from 0.92 to 0.99 for deletions and 0.91 to 0.92 for duplications.

Furthermore, the updated models effectively reduced the overall number of predicted CNV samples within the full CORIELL cohort. This therefore reduces the workload for human confirmation during the sample review process in our application, to a significantly smaller subset of predicted candidates. Initially, the preliminary deletion and duplication models predicted 925 and 333 samples with scores over 0.8, respectively. The updated deletion and duplication models reduced this to only 100 and 126 samples with predictions over 0.8. Among the 100 *PRKN* samples with predicted values exceeding 0.8 for deletions, 96 samples exhibited visible CNVs and were consequently integrated into the final deletion training set. Similarly, from the pool of 126 *PRKN* duplication candidates with prediction values of at least 0.8, the final duplication training set incorporated 53 samples, all of which attained prediction values exceeding 0.9. Supplementary Figure 3 showcases diverse, non-standard examples detected within CORIELL with predicted values of 1 to demonstrate the generalizability of our models.

These samples include homozygous deletions and noisy duplications which later enrich the feature spectrum captured by our training sets. Our goal with the updated training sets was to increase the overall accuracy and further reduce the Brier score (Supplementary Table 9) for both preliminary models.

### Final Model Performance

#### MLPA Validation on PRKN and SNCA

MLPA currently serves as the gold standard for confirming deletions and duplications in exonic regions that are crucial to determining disease relevance. Therefore it is imperative that we validate our predictions with this modality. We analyzed PFP and STELLENBOS samples in *PRKN*, our training set’s most prominent gene, and in *SNCA,* which was not included in model development. All samples with known mutations that could lead to false positive MLPA results were excluded. Although CNV-Finder can also detect intronic CNVs, these SVs were excluded from our comparisons to match the functionality of MLPA. Supplementary Table 11 shows our full MLPA validation results with additional performance listed for samples that are visible to our experts in LRR or BAF vs. position plots. Although validation on the full set shows strong performance with AUCs ranging from 0.83 to approximately 1, accuracy is even higher for visually identifiable samples, ranging from 0.92 to 1 due to the models’ dependence on features derived from these plots. Supplementary Figure 4 illustrates the concordance between NBA-based prediction values and visible plot features in *PRKN*.

We further confirmed that our duplication model captures both *SNCA* duplications and triplications with equally strong prediction values. Figure 3 features two monogenic families with duplications and triplications, demonstrating the generalizability of our models, despite the lack of any *SNCA* or triplication examples in the training sets. The lower frequency and clinical significance of CNVs in *SNCA* makes it crucial that our pipeline can find all positive samples without prior exposure to this gene. Moreover, the model maintained high specificity, assigning prediction values above 0.5 to less than 40 out of 1,234 samples, thereby limiting user review to only 3% of the dataset.

**Figure 3.**
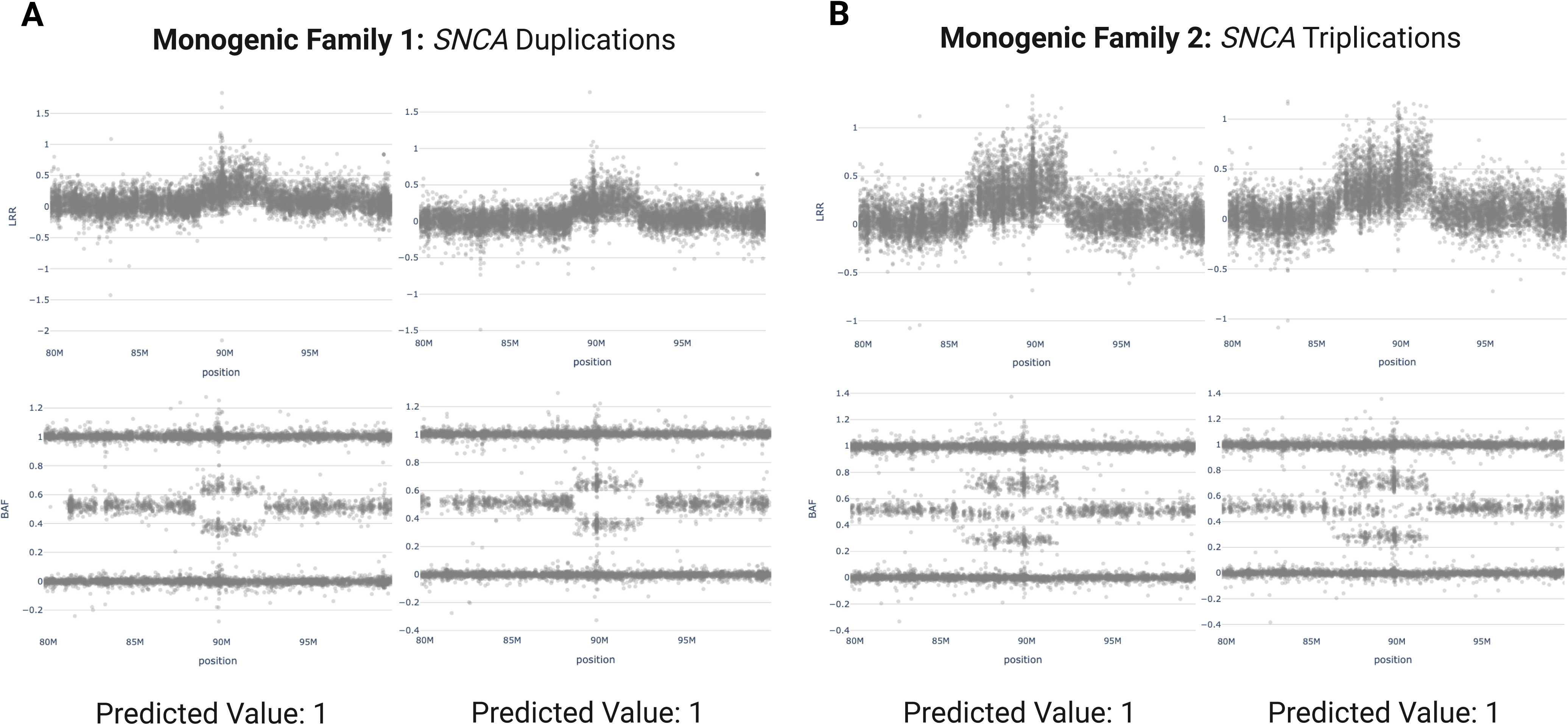
MLPA-validated *SNCA* multiplications from two monogenic families with their respective prediction values determined by CNV-Finder’s duplication model.

#### Short- and Long-Read Sequencing Validation

We used both Illumina short-read and ONT long-read WGS to validate predictions in the three genes incorporated in our model development. Long-read sequencing can identify distinct CNVs beyond the resolution of short reads and arrays, making it difficult to define false negatives in relation to array-based predictions ^2^. In contrast, short-read data is constrained by relatively high rates of false positives, also complicating the estimation of false negatives and potentially skewing true positive rates with respect to CNV-Finder’s results ^38^.

Our focus was on utilizing ONT long-read WGS data to validate samples with prediction values exceeding 0.9, to ensure that the strongest candidates from our models were not false positives. Figure 4 showcases five samples with predicted and confirmed *PRKN* deletions by long-read sequencing. These deletions vary in size, ranging from approximately 53 kb to 455 kb. Additionally, Figure 5 features three duplications near *MAPT* validated by long-read sequencing.

**Figure 4.**
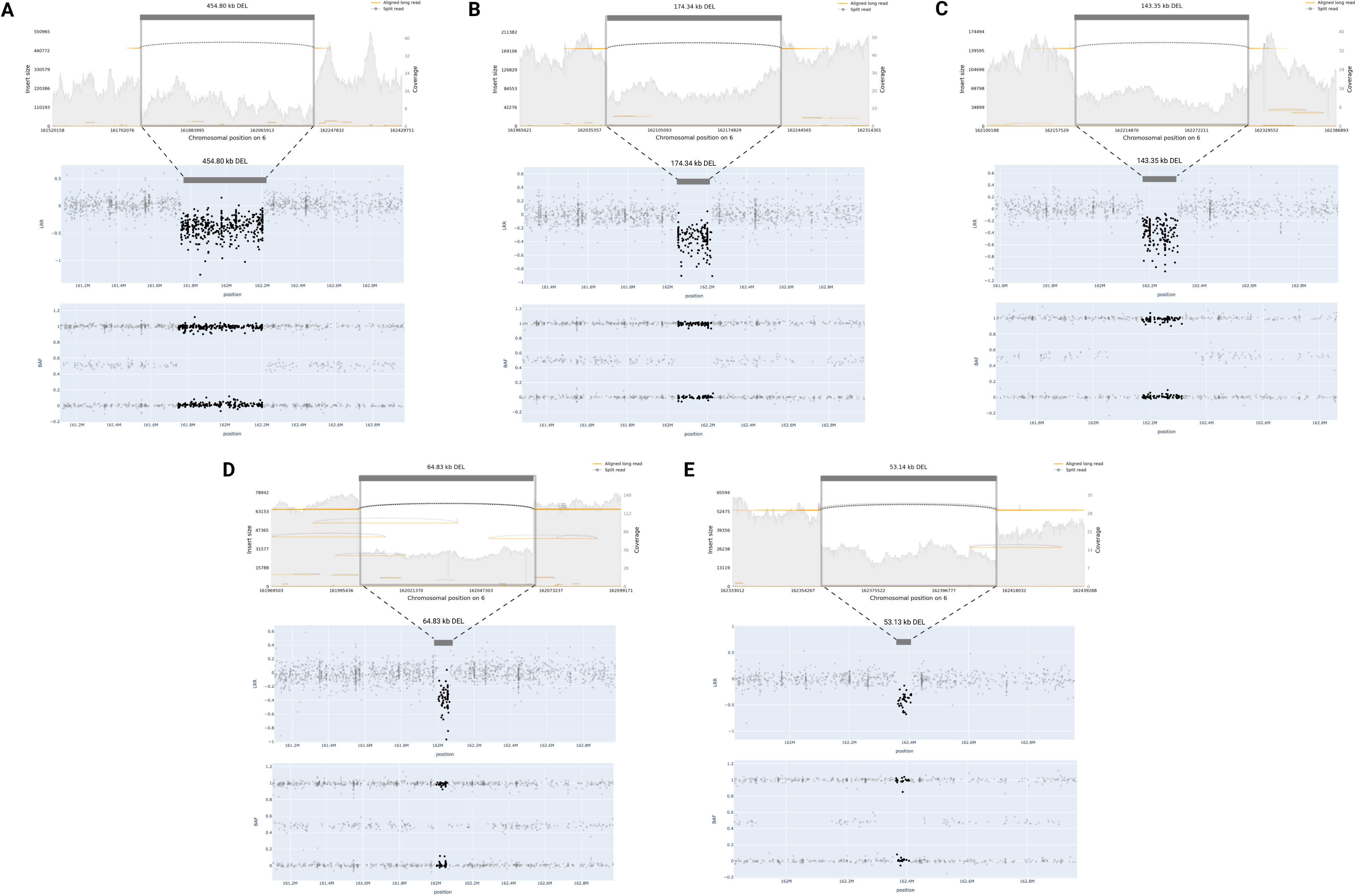
Long-read WGS validation of predicted deletions in *PRKN*. Four samples (A, B, C, E) received prediction values of 1 and one sample (D) received 0.93.

**Figure 5.**
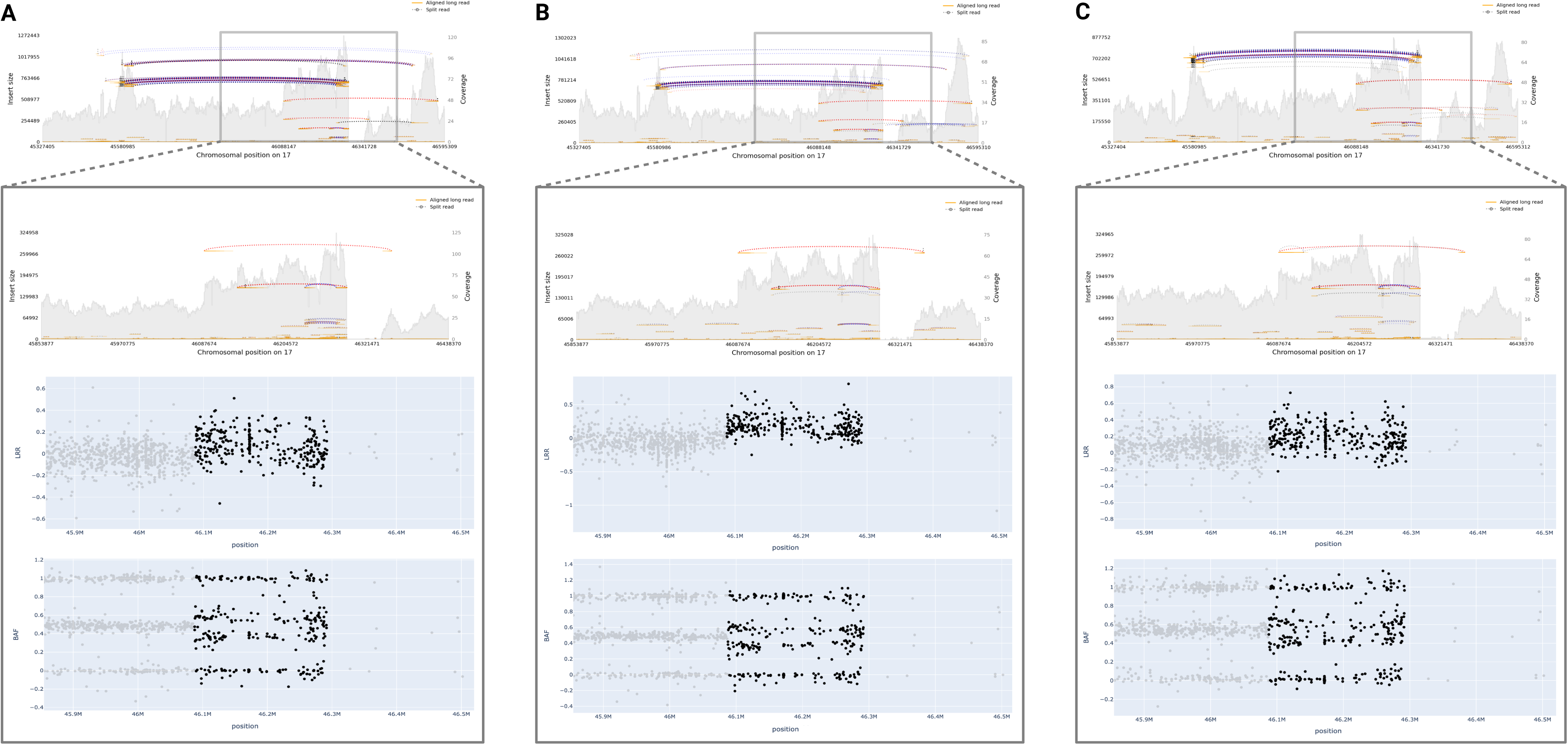
Long-read WGS validation of predicted duplications near *MAPT.* All three samples received prediction values of 1.

As previously mentioned, a 250 kb buffer surrounding the gene interval was employed in our analyses, enabling us to capture these duplications in the regions overlapping *MAPT* and *KANSL1*. Despite variations in the exact shape and variant noise levels, all validated CNVs received predicted values of 1.0 from their respective models, with only one predicted deletion obtaining a value of 0.93.

We incorporated short-read sequencing of an additional 337 samples to further explore predictions with values exceeding 0.5. Figure 6 presents CNVs confirmed by short-read WGS across a range of model prediction scores, with pink-highlighted regions indicating deletions or duplications identified by this modality. Model prediction values and visual clarity of the confirmed CNVs increase from left to right in the figure. Notably, *LINGO2* features the smallest deletion captured and confirmed by the model, with a width of only about 27 kb.

**Figure 6.**
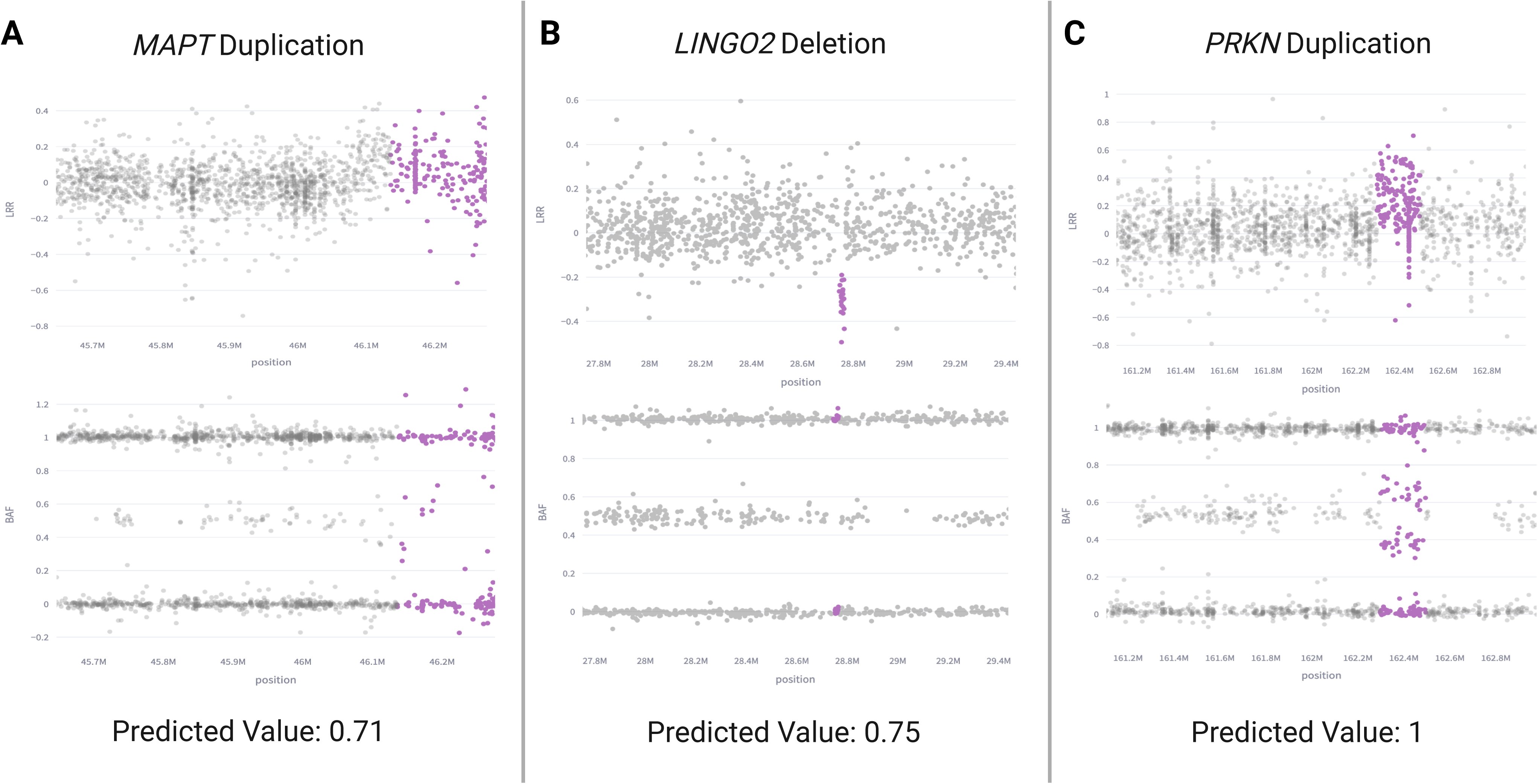
Short-read WGS validation of model results with varied prediction values.

Supplementary Table 12 breaks down the Positive Predictive Value (PPV) values for long-read and short-read WGS comparisons. PPV scores on samples with predicted values over 0.9 were high for long-read validation, demonstrating a score of 1 for *PRKN* deletions (*LINGO2* deletions were excluded due to insufficient samples) and 1 for duplications near *MAPT* (*PRKN* duplications were excluded due to insufficient samples). For short-read comparisons on samples with predicted values over 0.5, PPV averaged 0.95 for the final deletion model among *PRKN* and *LINGO2* while the final duplication model achieved an average of 0.90 for *PRKN* and *MAPT*. The leniency in *MAPT*’s false positive rate is attributed to the training set selections described in the following section.

#### Duplications within the 17q21.31 Region

Further exploration of the *MAPT* region highlights the crucial role of contextual understanding and human proficiency in enhancing the precision of CNV identification. Deliberate consideration was given to the classifications of “Yes,” “Maybe,” and “No” within the CNV-Finder application when selecting samples to add to the final training set. Supplementary Figure 5 showcases examples corresponding to each selection category. Samples D and E in this figure were marked as “Maybe” to avoid mislabeling them as negative samples with no CNVs. Typical duplication patterns are discernible in their LRR vs. position plots while the BAF vs. position plots lacked sufficient data. Additionally, potential artifacts like Sample F were marked as “Maybe” to prevent inclusion in the training sets. The duplication pattern highlighted in the “Yes” column, consistently appearing at locations around 461 kb to 463 kb on the 17th chromosome near *MAPT* and within the *KANSL1* region, exhibited the highest prevalence among the inspected CNVs across our three genes of interest. Across all cohorts reviewed for model development, a total of 737 samples exhibited visible duplications, while 414 samples showed no or limited evidence of duplication in the *MAPT* region, resulting in the annotation of 1151 samples with “Yes” or “No” for this variant.

As previously mentioned, the 17q21.31 region which encompasses the *MAPT* gene, is made complex by an H1 haplotype and inverted H2 haplotype. To better analyze this region and distinguish between haplotypes, we examined the H1/H2 tagging SNP rs8070723 across the 1151 annotated samples, revealing a Jaccard similarity score (Supplementary Table 9) of 0.94 between H2-carrier status (H1H2+H2H2) and the presence of an obvious duplication in the region (“Yes” classification). A logistic regression analysis was conducted on an 80:20 partition of these samples, ensuring consistency in the distributions of population ancestries across both data subsets. This resulted in a 97% accuracy and an AUC of 0.95 when predicting the existence of a duplication in this region using only H2-carrier status as the feature. There were only 6 false positives and 0 false negatives in the test set comprising 146 samples with the observed duplication and 85 samples without.

Furthermore, to better examine this variant in H2 non-carriers (H1H1), we applied our final duplication model to AD and dementia cases in the MDGAP-DEMENTIA and REGARDS cohorts. Among the 506 cases included in these cohorts, H2 non-carriers constitute 64%. Upon visual inspection of all samples with predicted values over 0.5, 38 instances were found to meet the criteria typically associated with a “Yes’’ designation in Supplementary Figure 5. These instances encompass a variety of NDDs, including cases of AD, FTD, PSP, DLB, MSA, CBD/CBS, Undetermined-Dementia, and other tauopathies; however, all are H2-carriers. Conversely, out of the 75 samples annotated as “Maybe” due to sparse BAF vs. chromosome position plots resembling those seen in plots D and E of Supplementary Figure 5, all are H2 non-carriers (H1H1). One sample reported as a “No Call” for SNP rs8070723 was categorized as “No.” Had the duplication model not been trained with careful consideration for the expansion of both binary classes (CNV exists vs. CNV does not exist), the final model would not have assigned high probabilities to H2 non-carriers, thus neglecting their flagging for human review. This highlights the significance of contextual interpretation by researchers and the additional layer of insight their expertise provides, particularly when array-based plots may not fully reflect all pertinent information.

### Utility

CNV-Finder facilitates streamlined and comprehensive analysis of deletions and duplications, capable of handling sample sizes necessary for capturing rare SVs and enhancing the statistical power of disease associations. This pipeline operates independently of outputs from any supplementary CNV prediction or imaging software, harnessing the extensive advantages of contemporary deep learning models. Additionally, its optional semi-automated functionality allows for the integration of expert validation, a process traditionally essential for confirming predicted candidates from standard array-based callers. Human expertise proves invaluable, particularly in complex regions like 17q21.31, where additional context is needed for accurate interpretation of detected variants or for navigating artifacts that may mimic structural variant features. Models trained with user visual confirmation have demonstrated enhanced detection capabilities for noisy and small CNVs, concurrently reducing false positive rates.

### Limitations and Next Steps

It is noteworthy that while human validation within CNV-Finder’s semi-automated approach has enhanced model performance in our analyses, it can also introduce inconsistencies and biases to predictions. The balance and representation of each class (samples with CNVs vs. samples without CNVs) in the training set may be influenced by selections in a manner detrimental to prediction accuracy.

In addition, our use cases have been confined to regional explorations with binary outputs from a fixed-length, sequence-to-vector LSTM model. This decision was made to facilitate rapid predictions on gene regions with established significance or interest demonstrated in the literature. Future iterations of the model will leverage LSTMs’ capacity to generate sequences of model prediction values. Although this adjustment may increase computational cost, it could enable users to apply the model across the entire genome, with outputs indicating more specific regions of elevated probability for each CNV type. We may also introduce new features that provide additional context to our predictions. These may involve a more selective inclusion of array probes in the input files, an addition of region-specific SNPs such as the *MAPT* H1/H2 tagging variant, or more granular classifications of CNV types, as in the case of *SNCA* triplications and duplications.

Future studies will need to investigate the performance across a wider spectrum of genotyping arrays, which may exhibit varying levels of call quality and coverage of genetic variants. Beyond this, the nature of our model architecture and its use of sliding windows, make it possible to transition our models to work on short-read WGS and potentially improve predictions through the enhancements in call quality from this modality.

Finally, as previously reported, the 17q21.31 region challenges the accuracy of short- and long-read SV calling ^29^. Conventional variant calling algorithms often struggle to precisely determine the position ranges of potential CNVs within this genomic region. For instance, Sniffles2, a widely-used open-source tool for analyzing long-read SVs, identified duplications in our PPMI samples proximal to *MAPT*, spanning positions 46,135,410 to 46,292,247 on chromosome 17, across both short-read and long-read sequencing data. However, upon visual examination of the three assessed samples (referenced in Figure 5) alongside SAMtools plots derived from long-read sequencing data, it became apparent that the start of the CNV likely occurred earlier than reported. Moreover, the loss of sequencing coverage immediately following the duplication complicates the accurate determination of the breakpoint. Regardless, our results align with prior research in this genomic locus, affirming the importance of investigating the genetic significance and phenotypic consequences of these identified duplication candidates ^39^.

## Conclusion

This study presents a comprehensive pipeline integrating deep learning techniques to expedite large-scale identification of CNVs within predefined genomic regions. Through application across five distinct use cases involving SVs within four genes implicated in NDD susceptibility, we showcase the adaptability and robustness of both our deletion and duplication models. By merging model predictions with domain-specific expertise, our approach significantly reduces the burden of manual validation, facilitating the identification of precise CNV candidates for subsequent downstream analyses or targeted validation using resource-intensive methods such as PCR or long-read sequencing.

## Data and Code Availability

This pipeline is implemented in Python and applied in Jupyter notebooks available for reference at: https://github.com/GP2code/CNV-Finder; DOI 10.5281/zenodo.14182563. Further documentation and example data is included in this repository with all pre-trained models saved in the CNV-Finder/ref_files/models folder.

## Acknowledgments, Funding, and COI

This research was supported in part by the Intramural Research Program of the NIH, National Institute on Aging (NIA), National Institutes of Health, Department of Health and Human Services; project number ZO1 AG000534, as well as the National Institute of Neurological Disorders and Stroke. The contributions of the NIH author(s) were made as part of their official duties as NIH federal employees, are in compliance with agency policy requirements, and are considered Works of the United States Government. However, the findings and conclusions presented in this paper are those of the author(s) and do not necessarily reflect the views of the NIH or the U.S. Department of Health and Human Services.

Data (DOI 10.5281/zenodo.10962119, release 7) used in the preparation of this article were obtained from the Global Parkinson’s Genetics Program (GP2). GP2 is funded by the Aligning Science Across Parkinson’s (ASAP) initiative and implemented by The Michael J. Fox Foundation for Parkinson’s Research (https://gp2.org). For a complete list of GP2 members see https://gp2.org.

N.K., M.B.M, M.J.K, K.L., H.L.L., D.V. and M.A.N.’s participation in this project was part of a competitive contract awarded to DataTecnica LLC by the National Institutes of Health to support open science research. M.A.N. also currently serves on the scientific advisory board for Character Bio Inc plus is a scientific founder at Neuron23 Inc and owns stock.

## Supplementary Information

**Supplementary Table 1.**
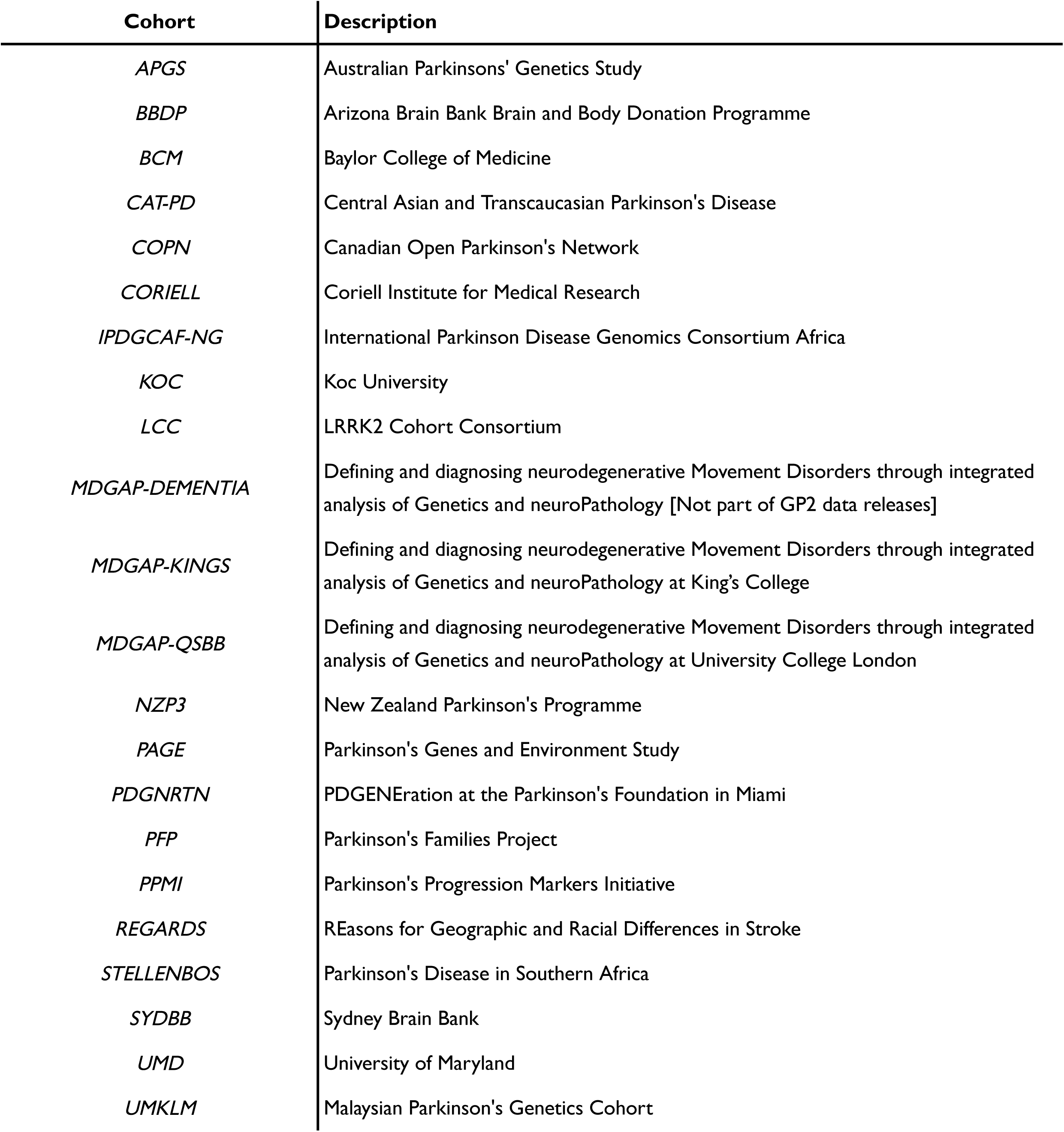
Descriptions of GP2 cohorts involved in training and validating our models.

**Supplementary Table 2.**
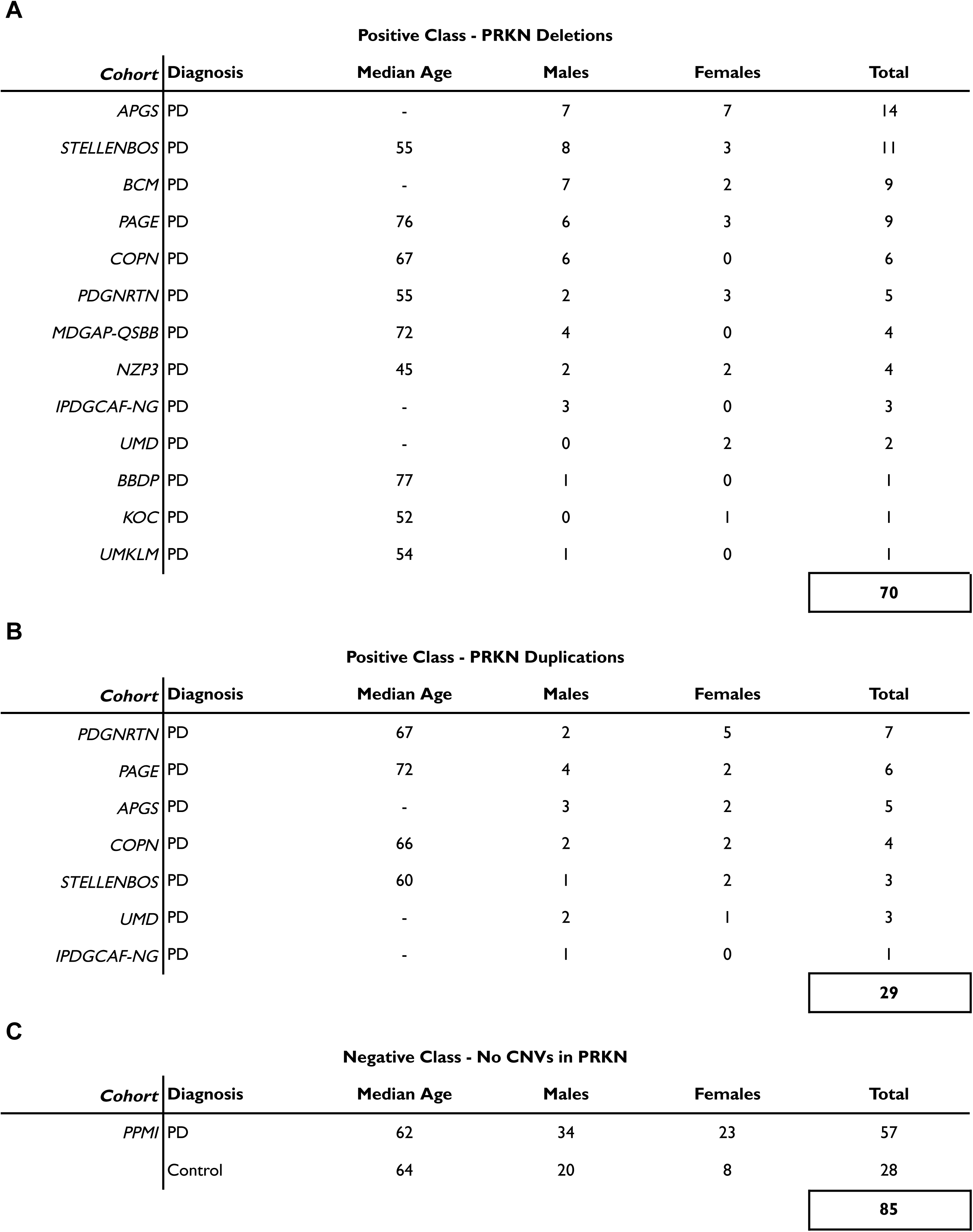
Summary of training set from expert annotations of *PRKN* interval. (A) *PRKN* deletions (B) *PRKN* duplications (C) Negative samples with no CNVs.

**Supplementary Table 3.**
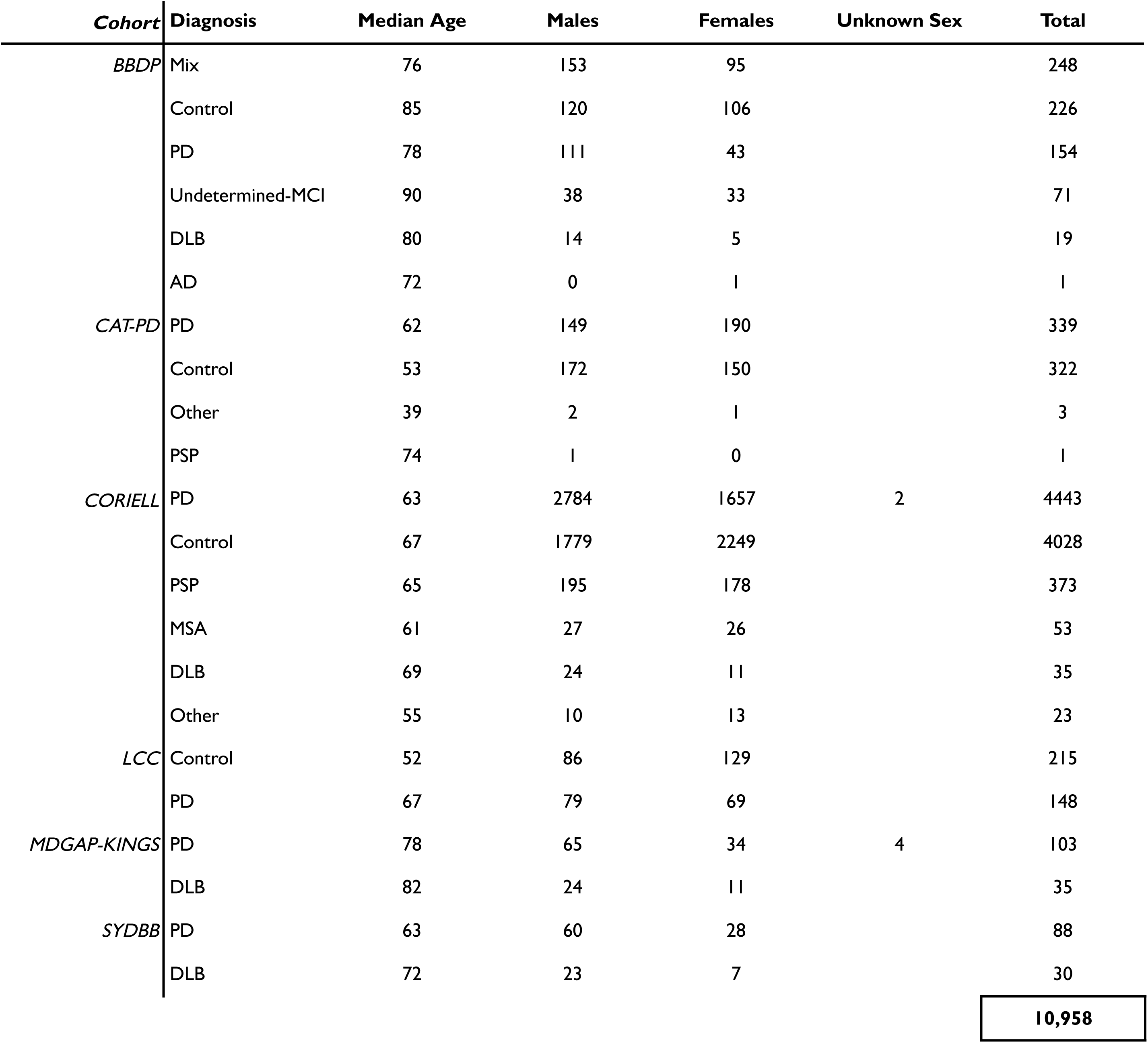
Summary statistics of GP2 cohorts used in training our updated and final models. “Undetermined-MCI” represents undiagnosed mild cognitive impairment and “Prodromal” represents prodromal PD with no motor symptoms. “MSA” refers to Multiple System Atrophy.

**Supplementary Table 4.**
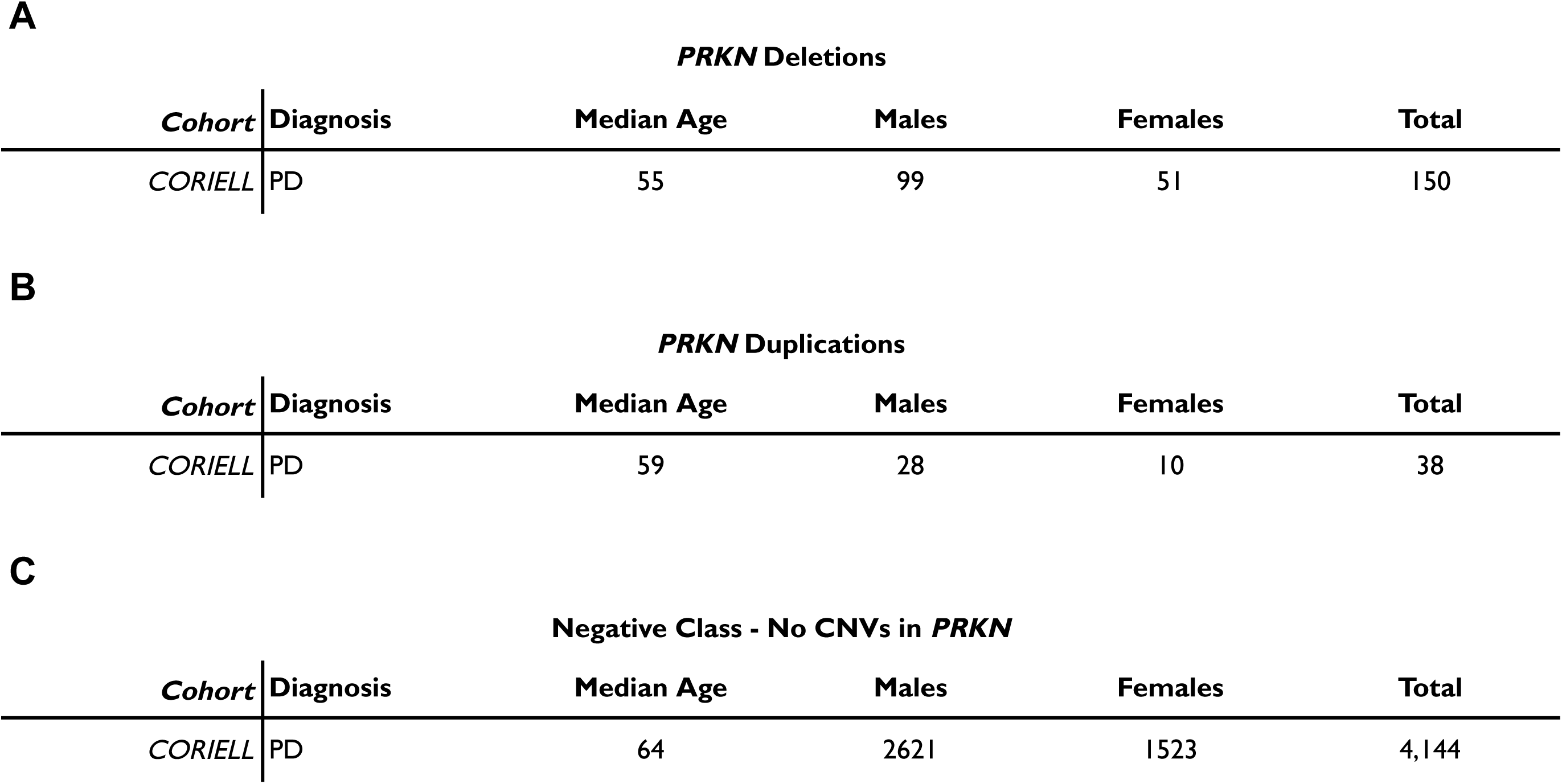
Summary statistics of held-out expert annotations for validation of preliminary and updated models on CORIELL samples. CORIELL samples were incorporated in the training set of our final models.

**Supplementary Table 5.**
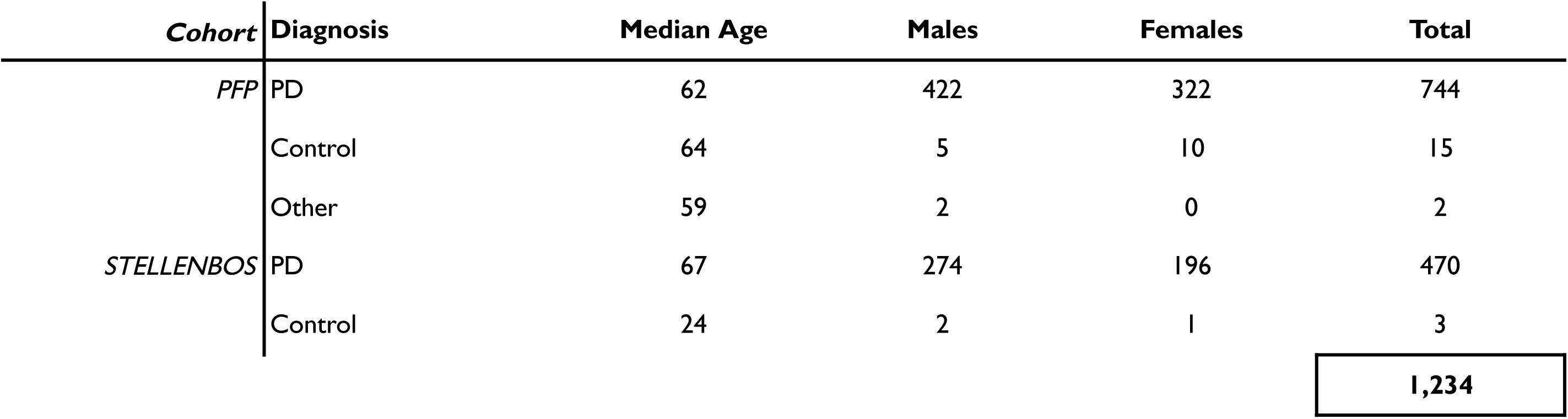
Summary statistics of PFP and STELLENBOS samples with MLPA data for prediction validation.

**Supplementary Table 6.**
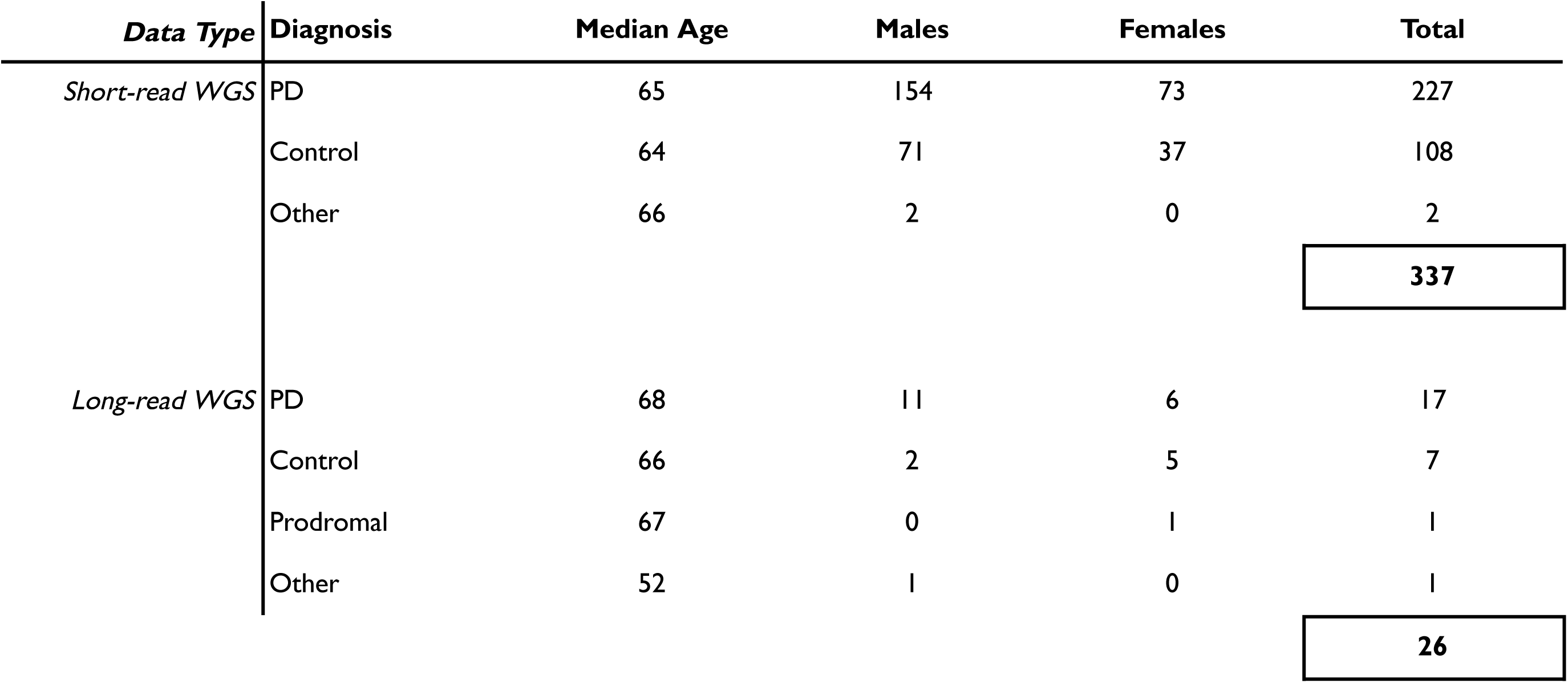
Summary of PPMI extension data available for prediction validation. No samples described here overlap with any PPMI samples incorporated in the models’ training sets.

**Supplementary Table 7.**
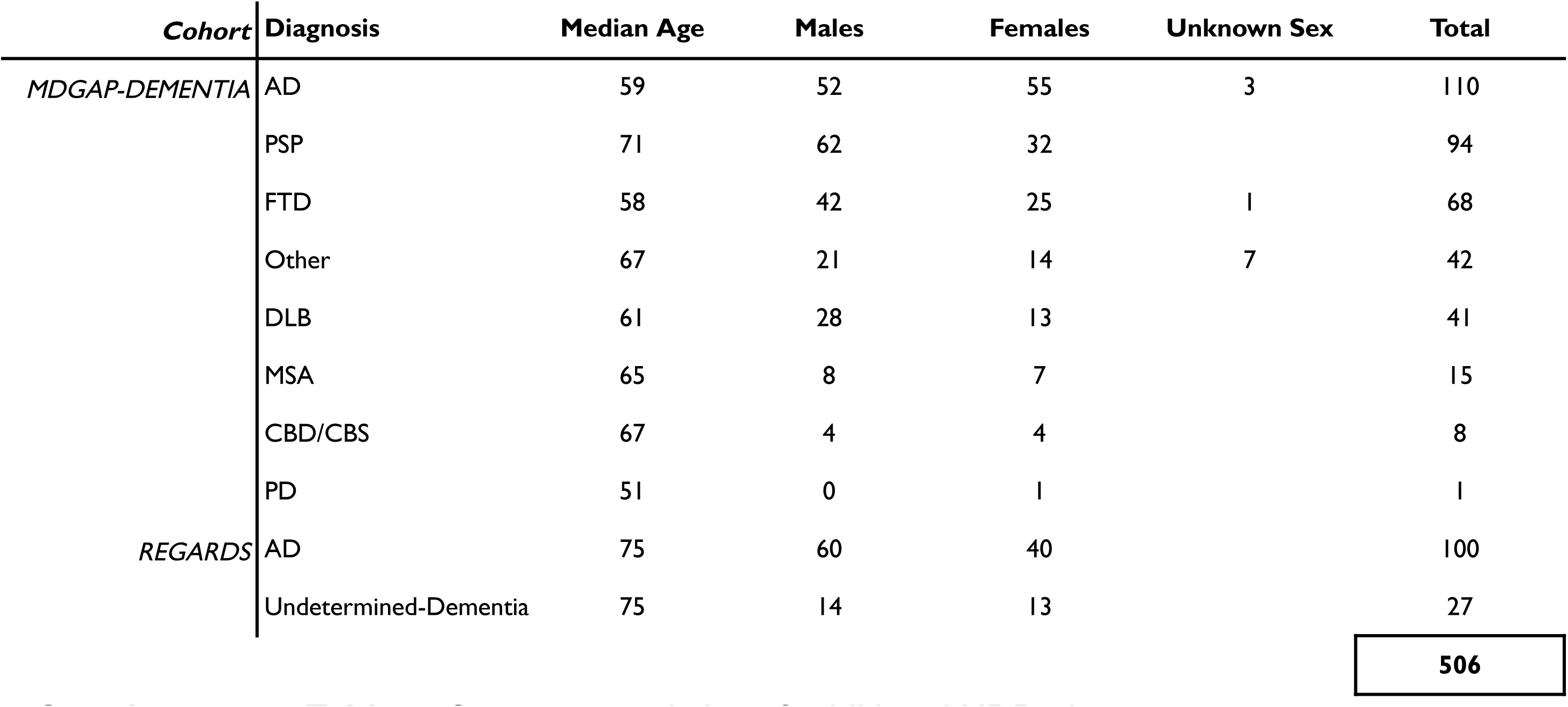
Summary statistics of additional NDD phenotype cases.

**Supplementary Figure 1.**
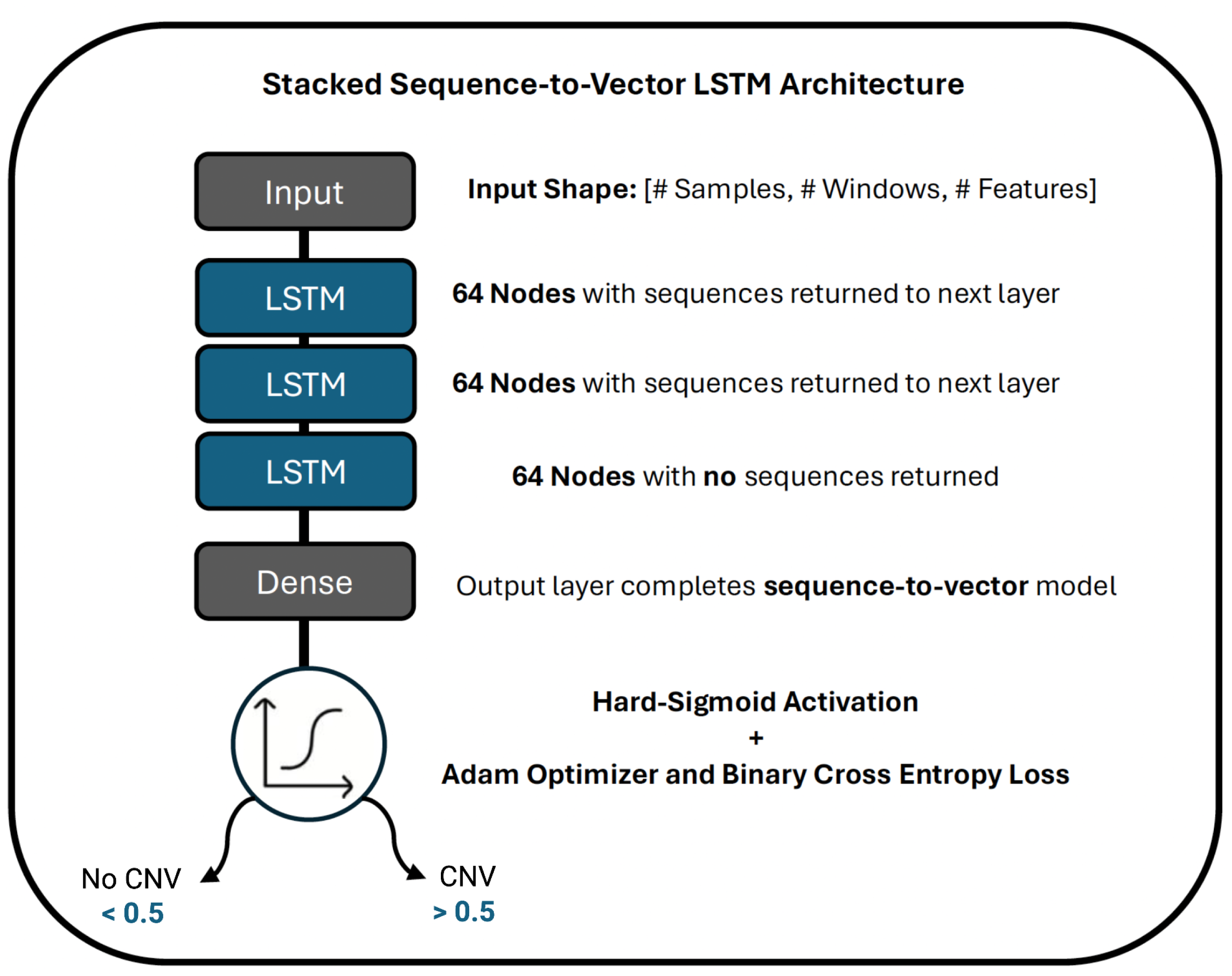
Overview of Long Short-Term Memory model architecture.

**Supplementary Figure 2.**
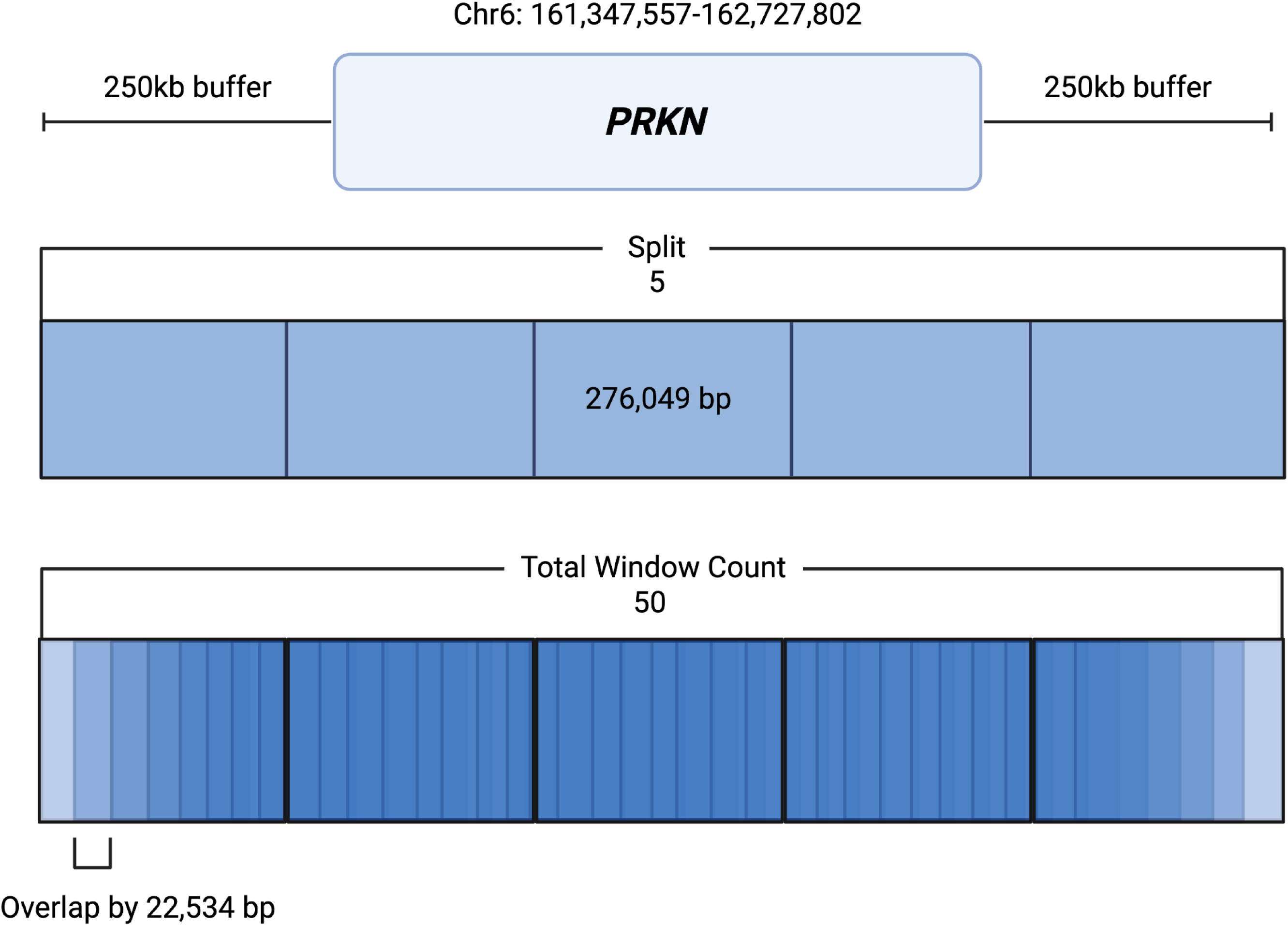
Overview of customizable “Split” and “Total Window Count” metrics to calculate sequential model features across gene intervals in base pairs (bp). The 250 kilobase (kb) buffer is an additional modifiable parameter.

**Supplementary Table 8.**
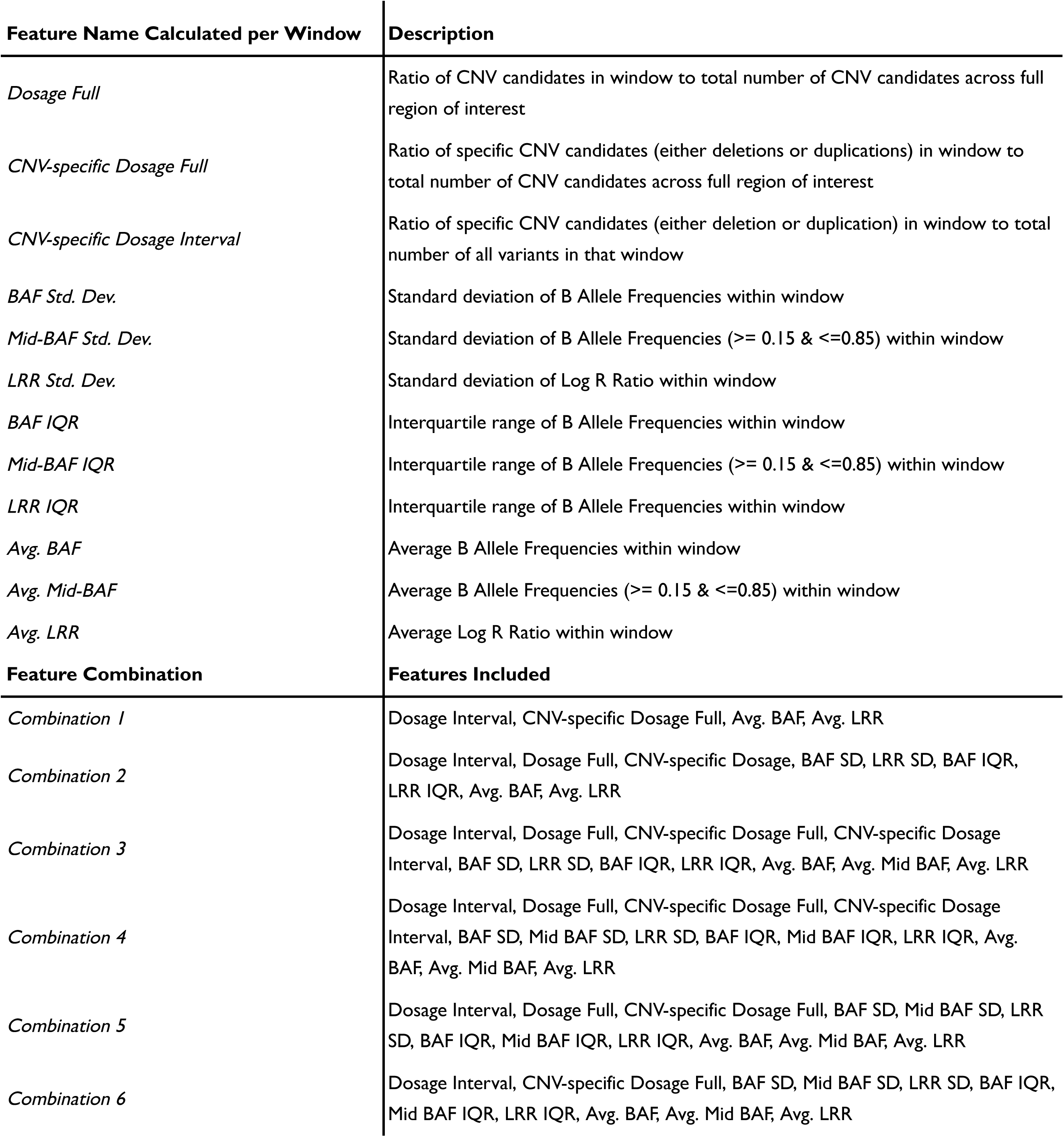
Descriptions of features and tested combinations used in model training. “Avg.” stands for average, “IQR” represents interquartile range, and “SD” refers to standard deviation.

**Supplementary Table 9.**
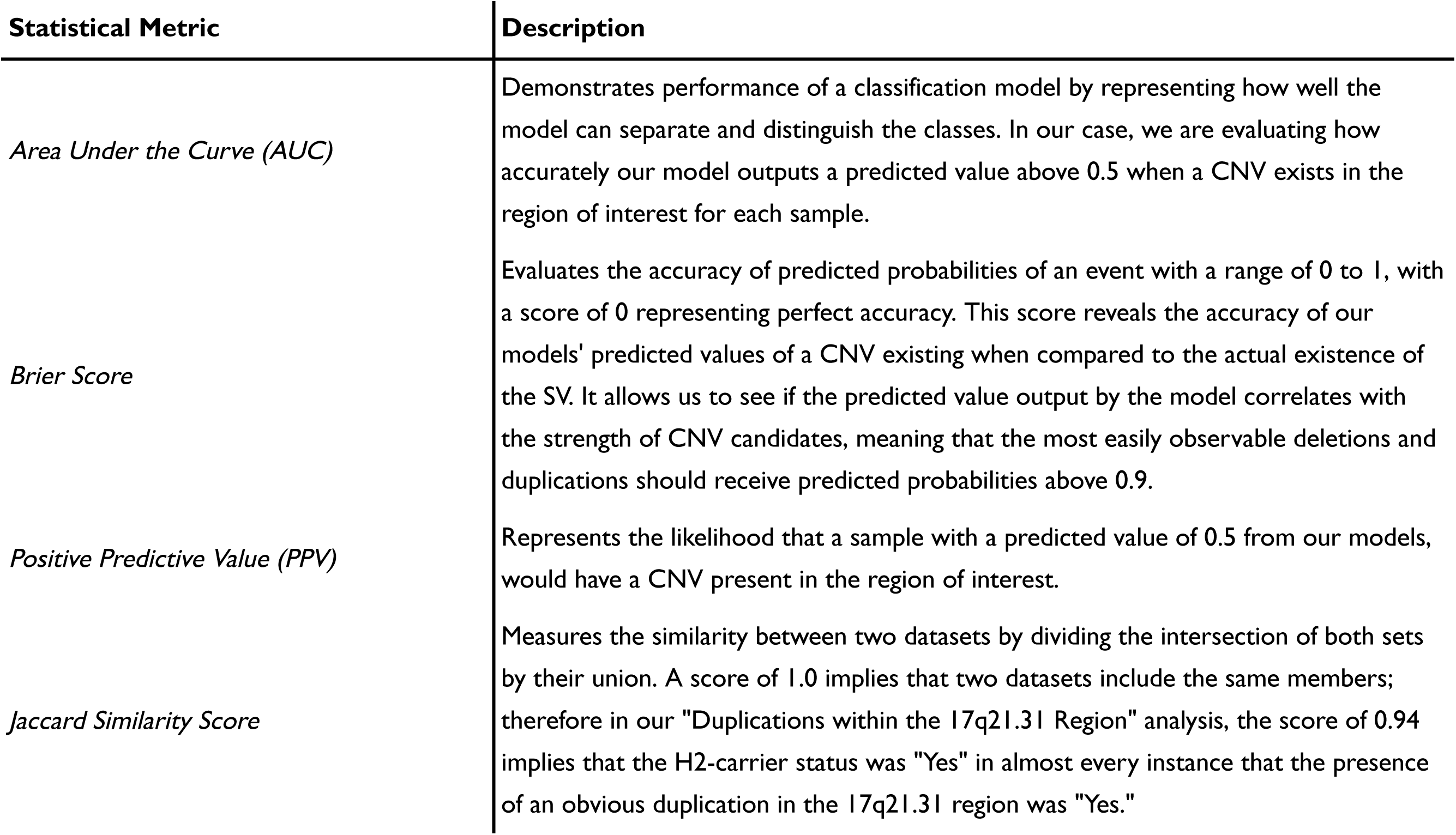
Descriptions of statistical metrics used in assessing model performance.

**Supplementary Table 10.**
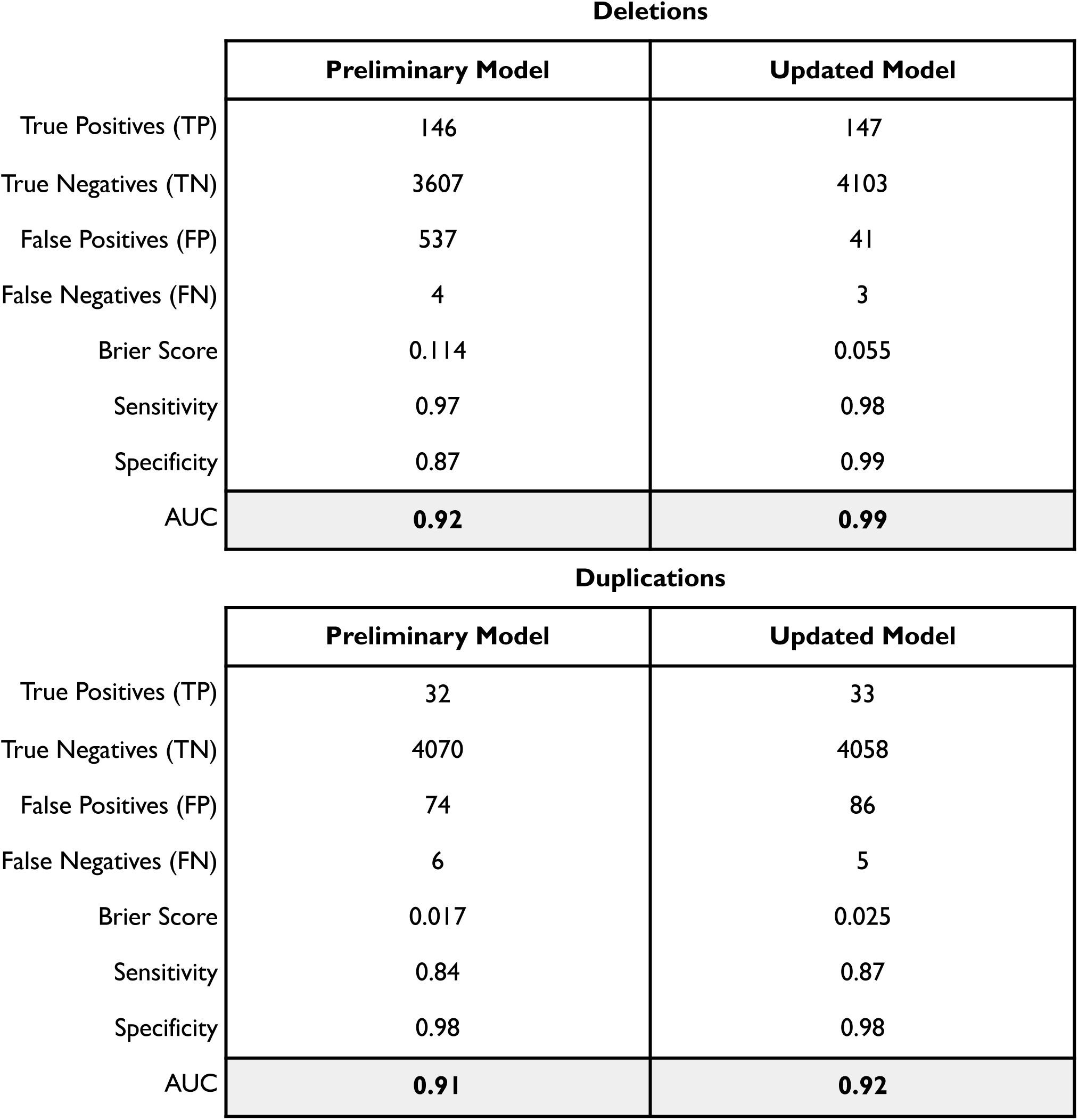
Preliminary and updated model results for known *PRKN* CNVs in held-out CORIELL PD Cases. Visually-confirmed samples with predicted values above 0.8 were fed back into the updated model’s training set to create the training sets for our final models.

**Supplementary Figure 3.**
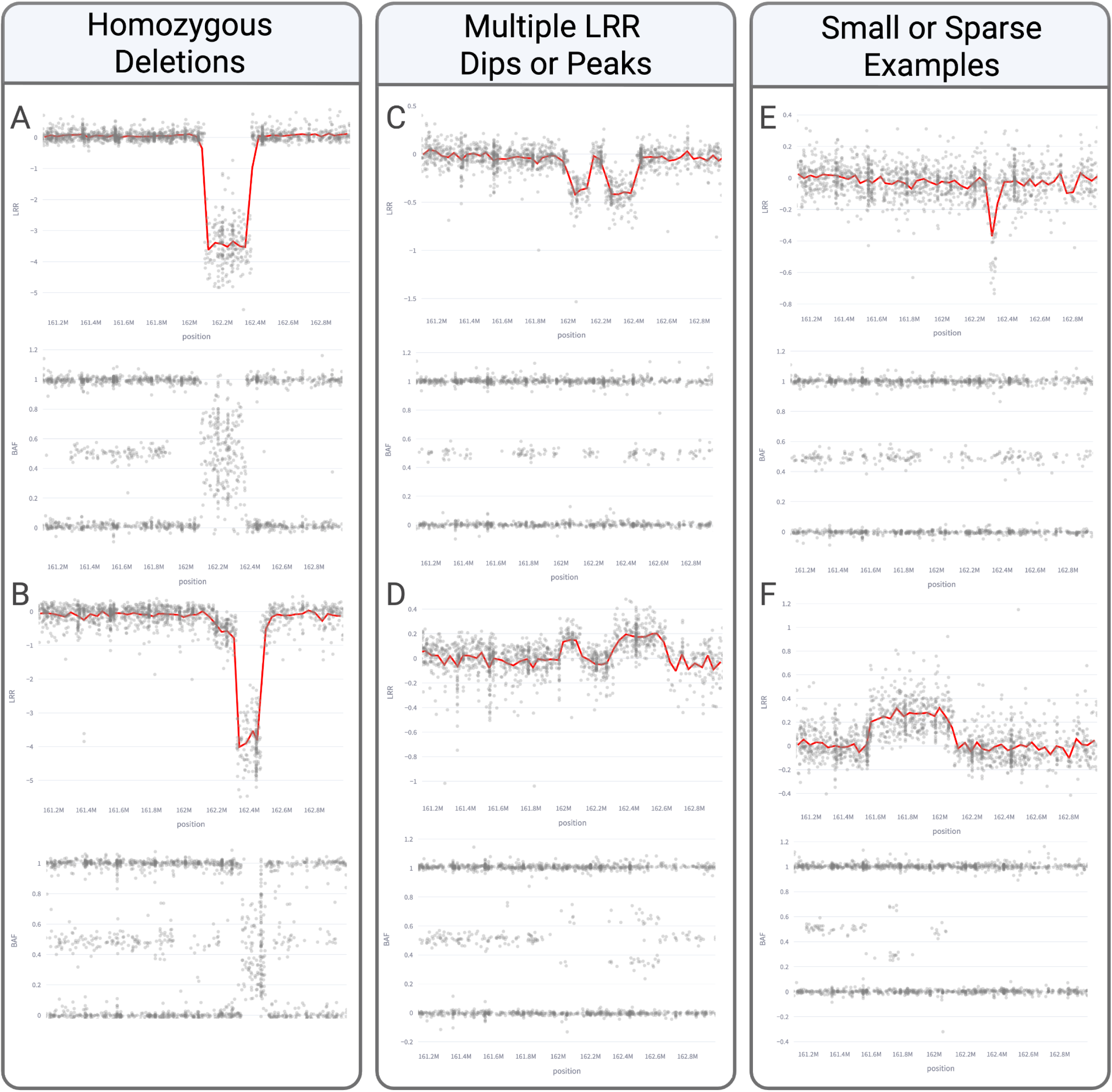
Atypical samples identified by models without explicit training of these categories.

**Supplementary Table 11.**
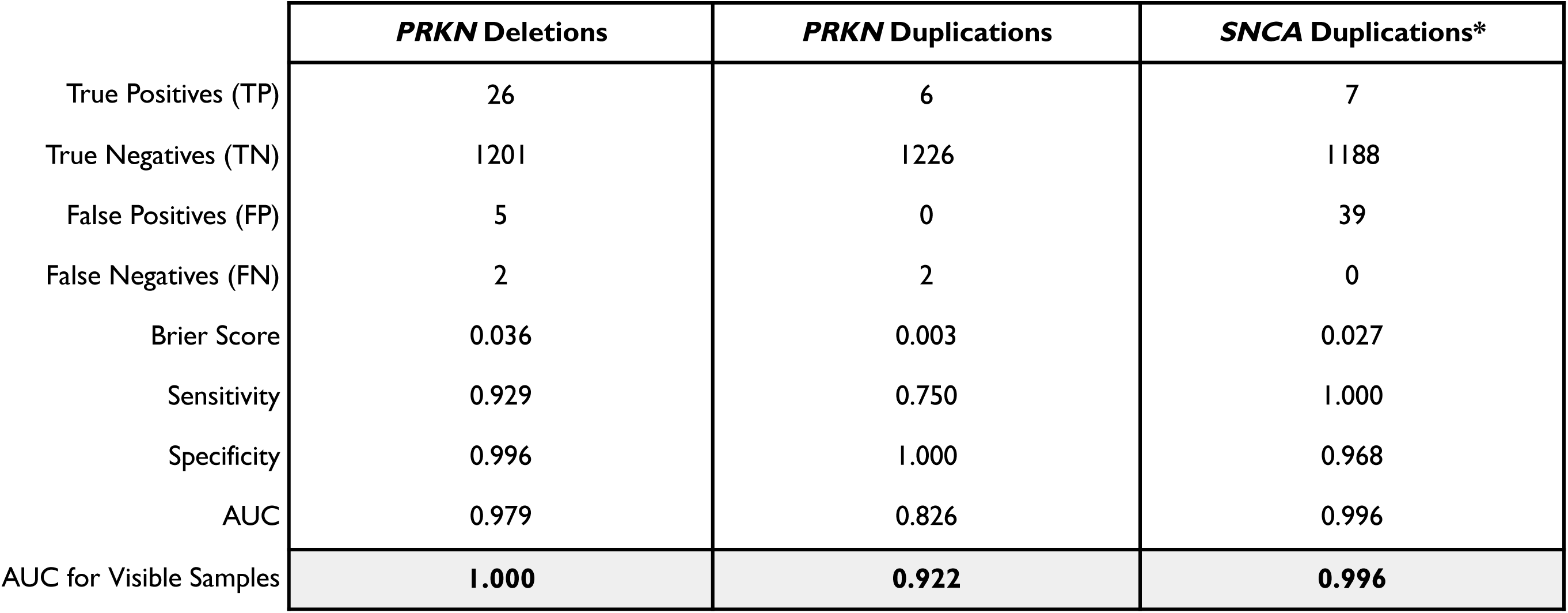
Summary of model predictions on combined MLPA data from PFP and STELLENBOS cohorts. Asterisks denotes that the final duplication model was used to make predictions on *SNCA;* however, both duplications and triplications were identified.

**Supplementary Figure 4.**
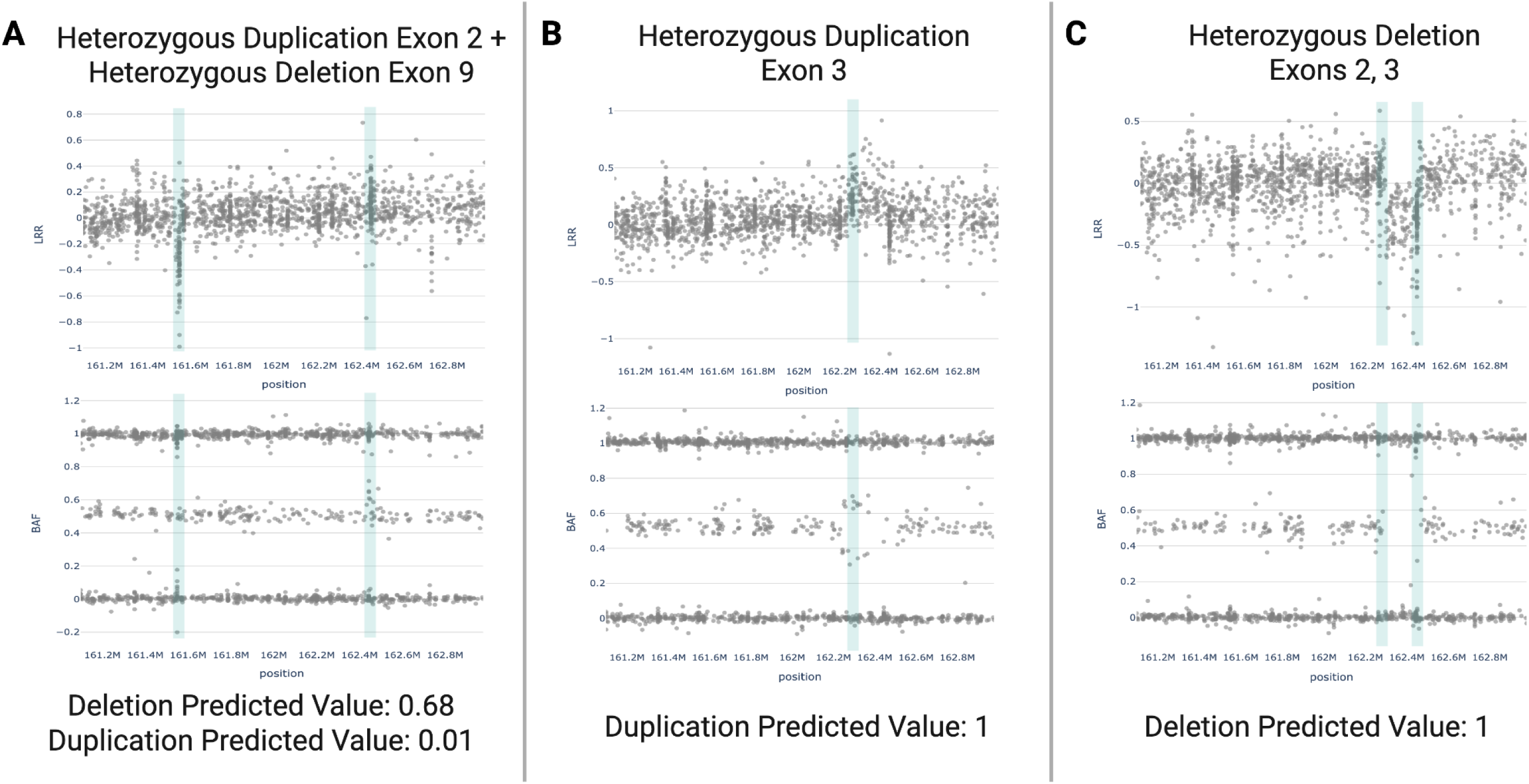
Samples with MLPA hits in *PRKN* and the comparison of NBA-based plot features to the models’ predicted values. Plots A and C feature Exon 2 which ranges from around 162,443,310-162,443,473 bp, Exon 9 in plot A spans approximately 161,548,854-161,549,003 bp, and Exon 3 in samples B and C includes positions 162,262,525-162,262,765 bp.

**Supplementary Table 12.**
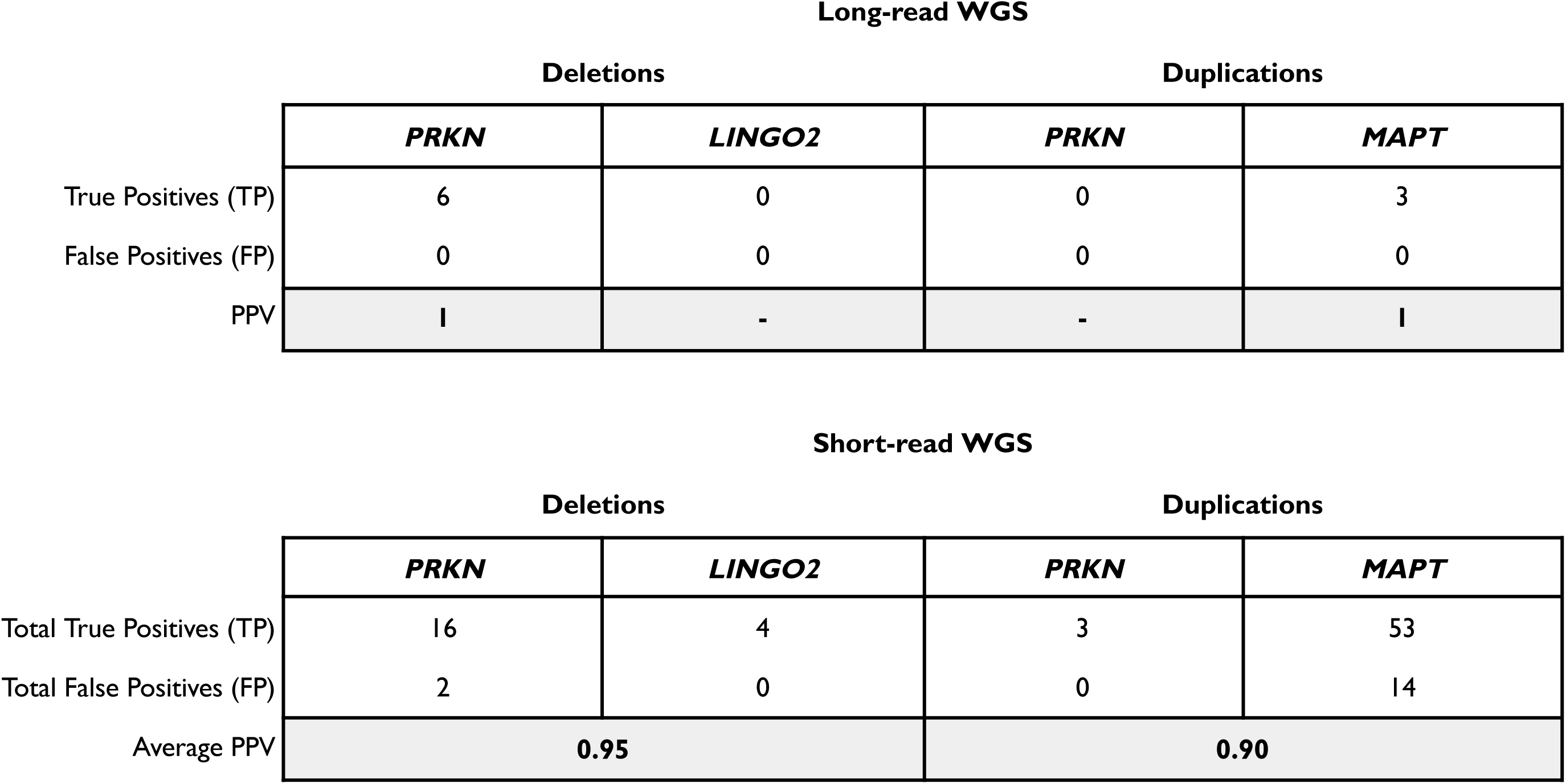
Summary of Positive Predictive Values (PPV) for final model predictions above 0.9 on PPMI long-read WGS data and predictions above 0.5 on short-read WGS data.

**Supplementary Figure 5.**
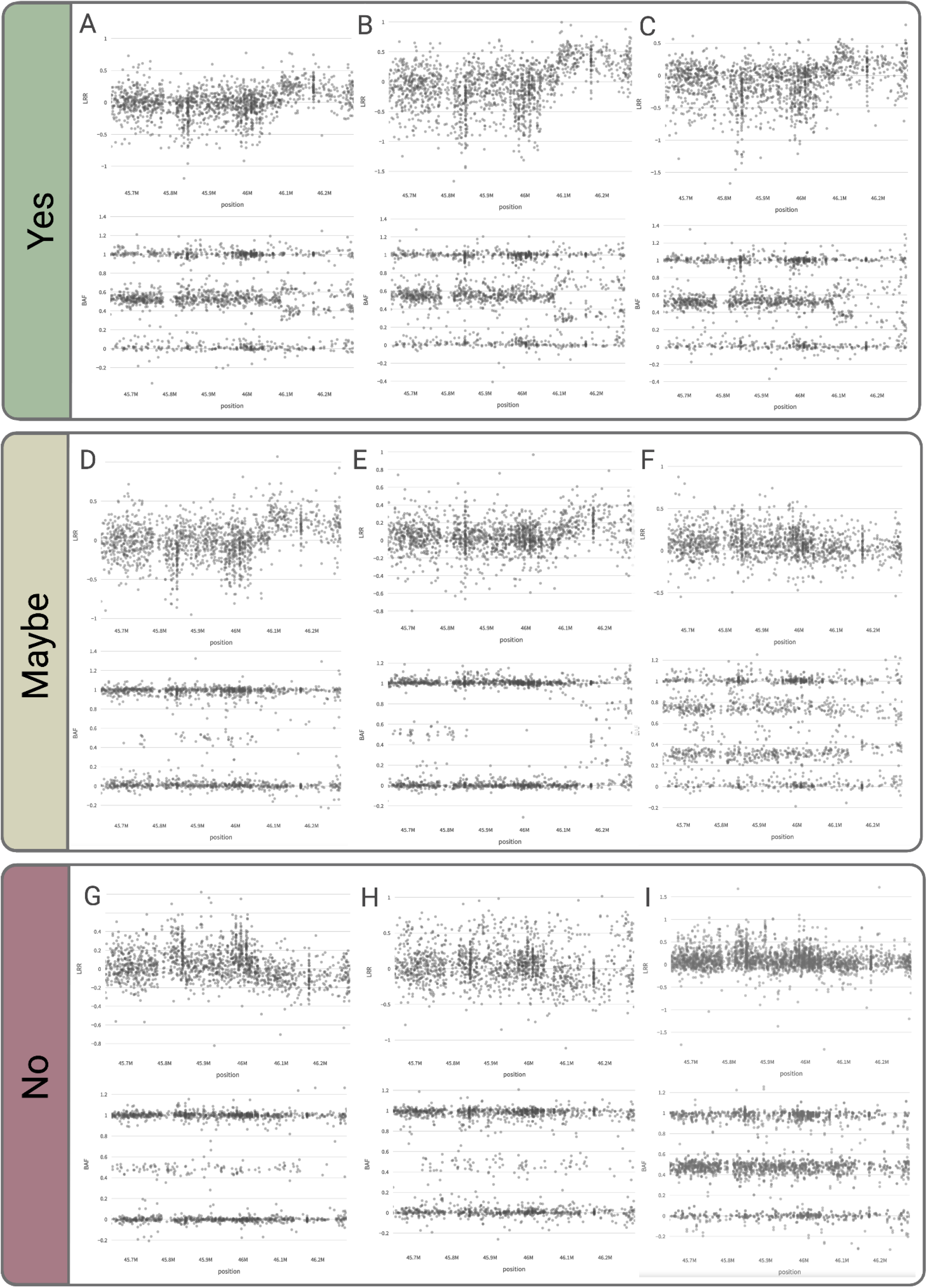
Examples of classification used on duplication near *MAPT* during visual confirmation.

