## supplementary_tables for "CNV-Finder: Streamlining Copy Number Variation Discovery"

| Cohort | Description |
|---|---|
| *APGS* | Australian Parkinsons' Genetics Study |
| *BBDP* | Arizona Brain Bank Brain and Body Donation Programme |
| *BCM* | Baylor College of Medicine |
| *CAT-PD* | Central Asian and Transcaucasian Parkinson's Disease |
| *COPN* | Canadian Open Parkinson's Network |
| *CORIELL* | Coriell Institute for Medical Research |
| *IPDGCAF-NG* | International Parkinson Disease Genomics Consortium Africa |
| *KOC* | Koc University |
| *LCC* | LRRK2 Cohort Consortium |
| *MDGAP-DEMENTIA* | Defining and diagnosing neurodegenerative Movement Disorders through integrated analysis of Genetics and neuroPathology [Not part of GP2 data releases] |
| *MDGAP-KINGS* | Defining and diagnosing neurodegenerative Movement Disorders through integrated analysis of Genetics and neuroPathology at King's College |
| *MDGAP-QSBB* | Defining and diagnosing neurodegenerative Movement Disorders through integrated analysis of Genetics and neuroPathology at University College London |
| *NZP3* | New Zealand Parkinson's Programme |
| *PAGE* | Parkinson's Genes and Environment Study |
| *PDGNRTN* | PDGENEration at the Parkinson's Foundation in Miami |
| *PFP* | Parkinson's Families Project |
| *PPMI* | Parkinson's Progression Markers Initiative |
| *REGARDS* | REasons for Geographic and Racial Differences in Stroke |
| *STELLENBOS* | Parkinson's Disease in Southern Africa |
| *SYDBB* | Sydney Brain Bank |
| *UMD* | University of Maryland |
| *UMKLM* | Malaysian Parkinson's Genetics Cohort |

**Supplementary Table 1**. Descriptions of GP2 cohorts involved in training and validating our models.

**A**

**Positive Class - PRKN Deletions**

| ***Cohort*** | **Diagnosis** | **Median Age** | **Males** | **Females** | **Total** |
|---|---|---|---|---|---|
| *APGS* | PD | - | 7 | 7 | 14 |
| *STELLENBOS* | PD | 55 | 8 | 3 | 11 |
| *BCM* | PD | - | 7 | 2 | 9 |
| *PAGE* | PD | 76 | 6 | 3 | 9 |
| *COPN* | PD | 67 | 6 | 0 | 6 |
| *PDGNRTN* | PD | 55 | 2 | 3 | 5 |
| *MDGAP-QSBB* | PD | 72 | 4 | 0 | 4 |
| *NZP3* | PD | 45 | 2 | 2 | 4 |
| *IPDGCAF-NG* | PD | - | 3 | 0 | 3 |
| *UMD* | PD | - | 0 | 2 | 2 |
| *BBDP* | PD | 77 | 1 | 0 | 1 |
| *KOC* | PD | 52 | 0 | 1 | 1 |
| *UMKLM* | PD | 54 | 1 | 0 | 1 |
| | | | | | **70** |

**B**

**Positive Class - PRKN Duplications**

| ***Cohort*** | **Diagnosis** | **Median Age** | **Males** | **Females** | **Total** |
|---|---|---|---|---|---|
| *PDGNRTN* | PD | 67 | 2 | 5 | 7 |
| *PAGE* | PD | 72 | 4 | 2 | 6 |
| *APGS* | PD | - | 3 | 2 | 5 |
| *COPN* | PD | 66 | 2 | 2 | 4 |
| *STELLENBOS* | PD | 60 | 1 | 2 | 3 |
| *UMD* | PD | - | 2 | 1 | 3 |
| *IPDGCAF-NG* | PD | - | 1 | 0 | 1 |
| | | | | | **29** |

**C**

**Negative Class - No CNVs in PRKN**

| ***Cohort*** | **Diagnosis** | **Median Age** | **Males** | **Females** | **Total** |
|---|---|---|---|---|---|
| *PPMI* | PD | 62 | 34 | 23 | 57 |
| | Control | 64 | 20 | 8 | 28 |
| | | | | | **85** |

**Supplementary Table 2**. Summary of training set from expert annotations of *PRKN* interval. (A) *PRKN* deletions (B) *PRKN* duplications (C) Negative samples with no CNVs.

| *Cohort* | Diagnosis | Median Age | Males | Females | Unknown Sex | Total |
|---|---|---|---|---|---|---|
| *BBDP* | Mix | 76 | 153 | 95 | | 248 |
| | Control | 85 | 120 | 106 | | 226 |
| | PD | 78 | 111 | 43 | | 154 |
| | Undetermined-MCI | 90 | 38 | 33 | | 71 |
| | DLB | 80 | 14 | 5 | | 19 |
| | AD | 72 | 0 | 1 | | 1 |
| *CAT-PD* | PD | 62 | 149 | 190 | | 339 |
| | Control | 53 | 172 | 150 | | 322 |
| | Other | 39 | 2 | 1 | | 3 |
| | PSP | 74 | 1 | 0 | | 1 |
| *CORIELL* | PD | 63 | 2784 | 1657 | 2 | 4443 |
| | Control | 67 | 1779 | 2249 | | 4028 |
| | PSP | 65 | 195 | 178 | | 373 |
| | MSA | 61 | 27 | 26 | | 53 |
| | DLB | 69 | 24 | 11 | | 35 |
| | Other | 55 | 10 | 13 | | 23 |
| *LCC* | Control | 52 | 86 | 129 | | 215 |
| | PD | 67 | 79 | 69 | | 148 |
| *MDGAP-KINGS* | PD | 78 | 65 | 34 | 4 | 103 |
| | DLB | 82 | 24 | 11 | | 35 |
| *SYDBB* | PD | 63 | 60 | 28 | | 88 |
| | DLB | 72 | 23 | 7 | | 30 |
| | | | | | | **10,958** |

**Supplementary Table 3**. Summary statistics of GP2 cohorts used in training our updated and final models. “Undetermined-MCI” represents undiagnosed mild cognitive impairment and “Prodromal” represents prodromal PD with no motor symptoms. “MSA” refers to Multiple System Atrophy.

**A**

***PRKN* Deletions**

| *Cohort* | Diagnosis | Median Age | Males | Females | Total |
|---|---|---|---|---|---|
| *CORIELL* | PD | 55 | 99 | 51 | 150 |

**B**

***PRKN* Duplications**

| *Cohort* | Diagnosis | Median Age | Males | Females | Total |
|---|---|---|---|---|---|
| *CORIELL* | PD | 59 | 28 | 10 | 38 |

**C**

**Negative Class - No CNVs in *PRKN***

| *Cohort* | Diagnosis | Median Age | Males | Females | Total |
|---|---|---|---|---|---|
| *CORIELL* | PD | 64 | 2621 | 1523 | 4,144 |

**Supplementary Table 4**. Summary statistics of held-out expert annotations for validation of preliminary and updated models on CORIELL samples. CORIELL samples were incorporated in the training set of our final models.

| *Cohort* | Diagnosis | Median Age | Males | Females | Total |
|---|---|---|---|---|---|
| *PFP* | PD | 62 | 422 | 322 | 744 |
| | Control | 64 | 5 | 10 | 15 |
| | Other | 59 | 2 | 0 | 2 |
| *STELLENBOS* | PD | 67 | 274 | 196 | 470 |
| | Control | 24 | 2 | 1 | 3 |
| | | | | | **1,234** |

**Supplementary Table 5**. Summary statistics of PFP and STELLENBOS samples with MLPA data for prediction validation.

| *Data Type* | Diagnosis | Median Age | Males | Females | Total |
|---|---|---|---|---|---|
| *Short-read WGS* | PD | 65 | 154 | 73 | 227 |
| | Control | 64 | 71 | 37 | 108 |
| | Other | 66 | 2 | 0 | 2 |
| | | | | | **337** |
| *Long-read WGS* | PD | 68 | 11 | 6 | 17 |
| | Control | 66 | 2 | 5 | 7 |
| | Prodromal | 67 | 0 | 1 | 1 |
| | Other | 52 | 1 | 0 | 1 |
| | | | | | **26** |

**Supplementary Table 6**. Summary of PPMI extension data available for prediction validation. No samples described here overlap with any PPMI samples incorporated in the models' training sets.

| *Cohort* | Diagnosis | Median Age | Males | Females | Unknown Sex | Total |
|---|---|---|---|---|---|---|
| *MDGAP-DEMENTIA* | AD | 59 | 52 | 55 | 3 | 110 |
| | PSP | 71 | 62 | 32 | | 94 |
| | FTD | 58 | 42 | 25 | 1 | 68 |
| | Other | 67 | 21 | 14 | 7 | 42 |
| | DLB | 61 | 28 | 13 | | 41 |
| | MSA | 65 | 8 | 7 | | 15 |
| | CBD/CBS | 67 | 4 | 4 | | 8 |
| | PD | 51 | 0 | 1 | | 1 |
| *REGARDS* | AD | 75 | 60 | 40 | | 100 |
| | Undetermined-Dementia | 75 | 14 | 13 | | 27 |
| | | | | | | **506** |

**Supplementary Table 7**. Summary statistics of additional NDD phenotype cases.

| Feature Name Calculated per Window | Description |
|---|---|
| *Dosage Full* | Ratio of CNV candidates in window to total number of CNV candidates across full region of interest |
| *CNV-specific Dosage Full* | Ratio of specific CNV candidates (either deletions or duplications) in window to total number of CNV candidates across full region of interest |
| *CNV-specific Dosage Interval* | Ratio of specific CNV candidates (either deletion or duplication) in window to total number of all variants in that window |
| *BAF Std. Dev.* | Standard deviation of B Allele Frequencies within window |
| *Mid-BAF Std. Dev.* | Standard deviation of B Allele Frequencies (>= 0.15 & <=0.85) within window |
| *LRR Std. Dev.* | Standard deviation of Log R Ratio within window |
| *BAF IQR* | Interquartile range of B Allele Frequencies within window |
| *Mid-BAF IQR* | Interquartile range of B Allele Frequencies (>= 0.15 & <=0.85) within window |
| *LRR IQR* | Interquartile range of B Allele Frequencies within window |
| *Avg. BAF* | Average B Allele Frequencies within window |
| *Avg. Mid-BAF* | Average B Allele Frequencies (>= 0.15 & <=0.85) within window |
| *Avg. LRR* | Average Log R Ratio within window |
| **Feature Combination** | **Features Included** |
| *Combination 1* | Dosage Interval, CNV-specific Dosage Full, Avg. BAF, Avg. LRR |
| *Combination 2* | Dosage Interval, Dosage Full, CNV-specific Dosage, BAF SD, LRR SD, BAF IQR, LRR IQR, Avg. BAF, Avg. LRR |
| *Combination 3* | Dosage Interval, Dosage Full, CNV-specific Dosage Full, CNV-specific Dosage Interval, BAF SD, LRR SD, BAF IQR, LRR IQR, Avg. BAF, Avg. Mid BAF, Avg. LRR |
| *Combination 4* | Dosage Interval, Dosage Full, CNV-specific Dosage Full, CNV-specific Dosage Interval, BAF SD, Mid BAF SD, LRR SD, BAF IQR, Mid BAF IQR, LRR IQR, Avg. BAF, Avg. Mid BAF, Avg. LRR |
| *Combination 5* | Dosage Interval, Dosage Full, CNV-specific Dosage Full, BAF SD, Mid BAF SD, LRR SD, BAF IQR, Mid BAF IQR, LRR IQR, Avg. BAF, Avg. Mid BAF, Avg. LRR |
| *Combination 6* | Dosage Interval, CNV-specific Dosage Full, BAF SD, Mid BAF SD, LRR SD, BAF IQR, Mid BAF IQR, LRR IQR, Avg. BAF, Avg. Mid BAF, Avg. LRR |

**Supplementary Table 8**. Descriptions of features and tested combinations used in model training. "Avg." stands for average, "IQR" represents interquartile range, and "SD" refers to standard deviation.

| Statistical Metric | Description |
|---|---|
| *Area Under the Curve (AUC)* | Demonstrates performance of a classification model by representing how well the model can separate and distinguish the classes. In our case, we are evaluating how accurately our model outputs a predicted value above 0.5 when a CNV exists in the region of interest for each sample. |
| *Brier Score* | Evaluates the accuracy of predicted probabilities of an event with a range of 0 to 1, with a score of 0 representing perfect accuracy. This score reveals the accuracy of our models' predicted values of a CNV existing when compared to the actual existence of the SV. It allows us to see if the predicted value output by the model correlates with the strength of CNV candidates, meaning that the most easily observable deletions and duplications should receive predicted probabilities above 0.9. |
| *Positive Predictive Value (PPV)* | Represents the likelihood that a sample with a predicted value of 0.5 from our models, would have a CNV present in the region of interest. |
| *Jaccard Similarity Score* | Measures the similarity between two datasets by dividing the intersection of both sets by their union. A score of 1.0 implies that two datasets include the same members; therefore in our "Duplications within the 17q21.31 Region" analysis, the score of 0.94 implies that the H2-carrier status was "Yes" in almost every instance that the presence of an obvious duplication in the 17q21.31 region was "Yes." |

**Supplementary Table 9**. Descriptions of statistical metrics used in assessing model performance.

**Deletions**

| | Preliminary Model | Updated Model |
|---|---|---|
| True Positives (TP) | 146 | 147 |
| True Negatives (TN) | 3607 | 4103 |
| False Positives (FP) | 537 | 41 |
| False Negatives (FN) | 4 | 3 |
| Brier Score | 0.114 | 0.055 |
| Sensitivity | 0.97 | 0.98 |
| Specificity | 0.87 | 0.99 |
| AUC | **0.92** | **0.99** |

**Duplications**

| | Preliminary Model | Updated Model |
|---|---|---|
| True Positives (TP) | 32 | 33 |
| True Negatives (TN) | 4070 | 4058 |
| False Positives (FP) | 74 | 86 |
| False Negatives (FN) | 6 | 5 |
| Brier Score | 0.017 | 0.025 |
| Sensitivity | 0.84 | 0.87 |
| Specificity | 0.98 | 0.98 |
| AUC | **0.91** | **0.92** |

**Supplementary Table 10**. Preliminary and updated model results for known *PRKN* CNVs in held-out CORIELL PD Cases. Visually-confirmed samples with predicted values above 0.8 were fed back into the updated model's training set to create the training sets for our final models.

| | *PRKN* Deletions | *PRKN* Duplications | *SNCA* Duplications* |
|---|---|---|---|
| True Positives (TP) | 26 | 6 | 7 |
| True Negatives (TN) | 1201 | 1226 | 1188 |
| False Positives (FP) | 5 | 0 | 39 |
| False Negatives (FN) | 2 | 2 | 0 |
| Brier Score | 0.036 | 0.003 | 0.027 |
| Sensitivity | 0.929 | 0.750 | 1.000 |
| Specificity | 0.996 | 1.000 | 0.968 |
| AUC | 0.979 | 0.826 | 0.996 |
| AUC for Visible Samples | **1.000** | **0.922** | **0.996** |

**Supplementary Table 11**. Summary of model predictions on combined MLPA data from PFP and STELLENBOS cohorts. Asterisks denotes that the final duplication model was used to make predictions on *SNCA;* however, both duplications and triplications were identified.

**Long-read WGS**

| | Deletions | | Duplications | |
|---|---|---|---|---|
| | ***PRKN*** | ***LINGO2*** | ***PRKN*** | ***MAPT*** |
| True Positives (TP) | 6 | 0 | 0 | 3 |
| False Positives (FP) | 0 | 0 | 0 | 0 |
| PPV | **1** | **-** | **-** | **1** |

**Short-read WGS**

| | Deletions | | Duplications | |
|---|---|---|---|---|
| | ***PRKN*** | ***LINGO2*** | ***PRKN*** | ***MAPT*** |
| Total True Positives (TP) | 16 | 4 | 3 | 53 |
| Total False Positives (FP) | 2 | 0 | 0 | 14 |
| Average PPV | **0.95** | | **0.90** | |

**Supplementary Table 12**. Summary of Positive Predictive Values (PPV) for final model predictions above 0.9 on PPMI long-read WGS data and predictions above 0.5 on short-read WGS data.
