## supplementary_figures for "CNV-Finder: Streamlining Copy Number Variation Discovery"

### Supplementary Information

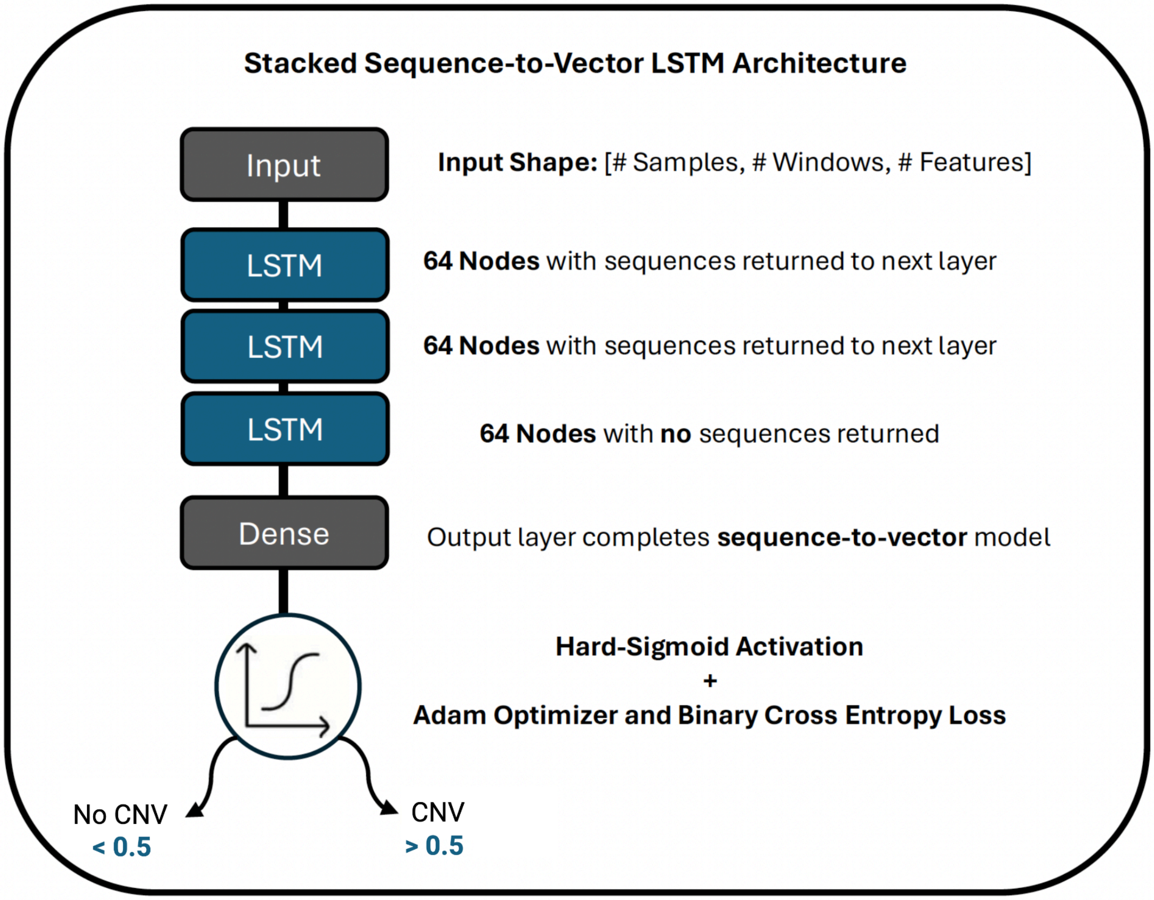


**Supplementary Figure 1**. Overview of Long Short-Term Memory model architecture.

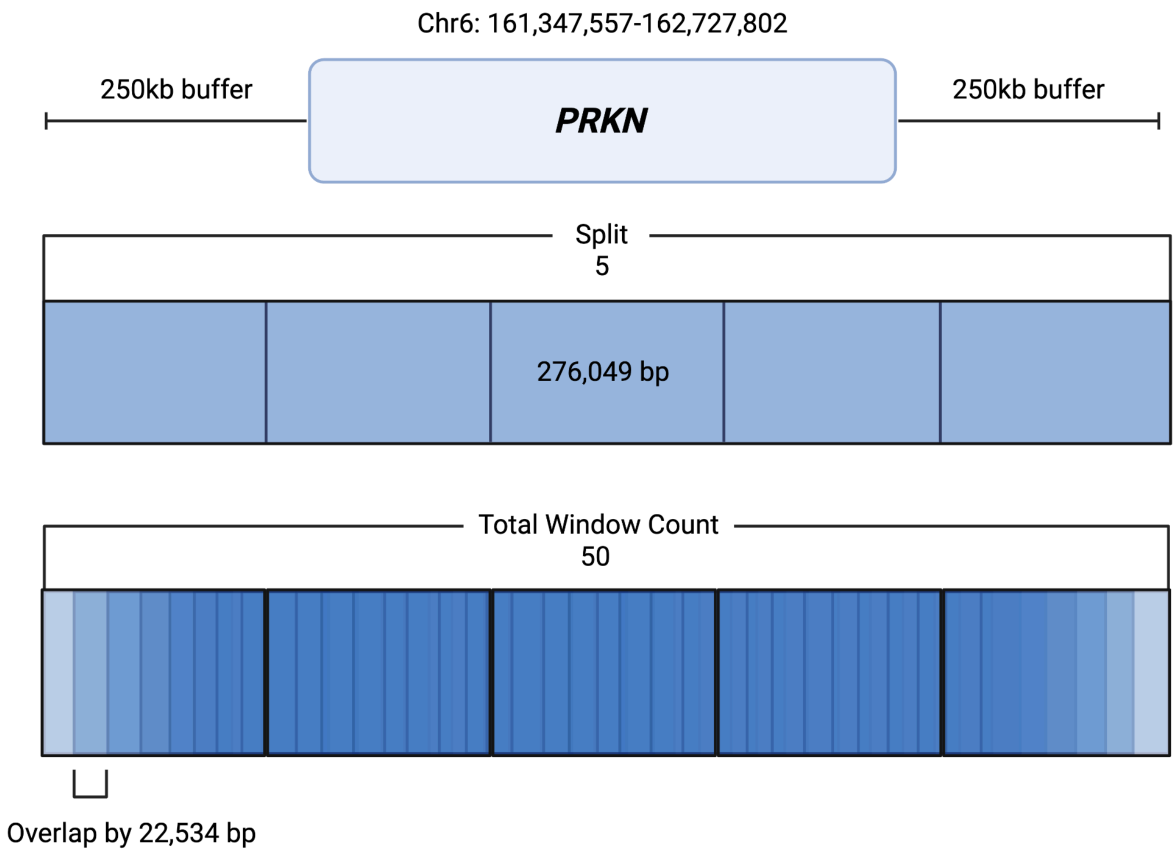


**Supplementary Figure 2**. Overview of customizable “Split” and “Total Window Count” metrics to calculate sequential model features across gene intervals in base pairs (bp). The 250 kilobase (kb) buffer is an additional modifiable parameter.

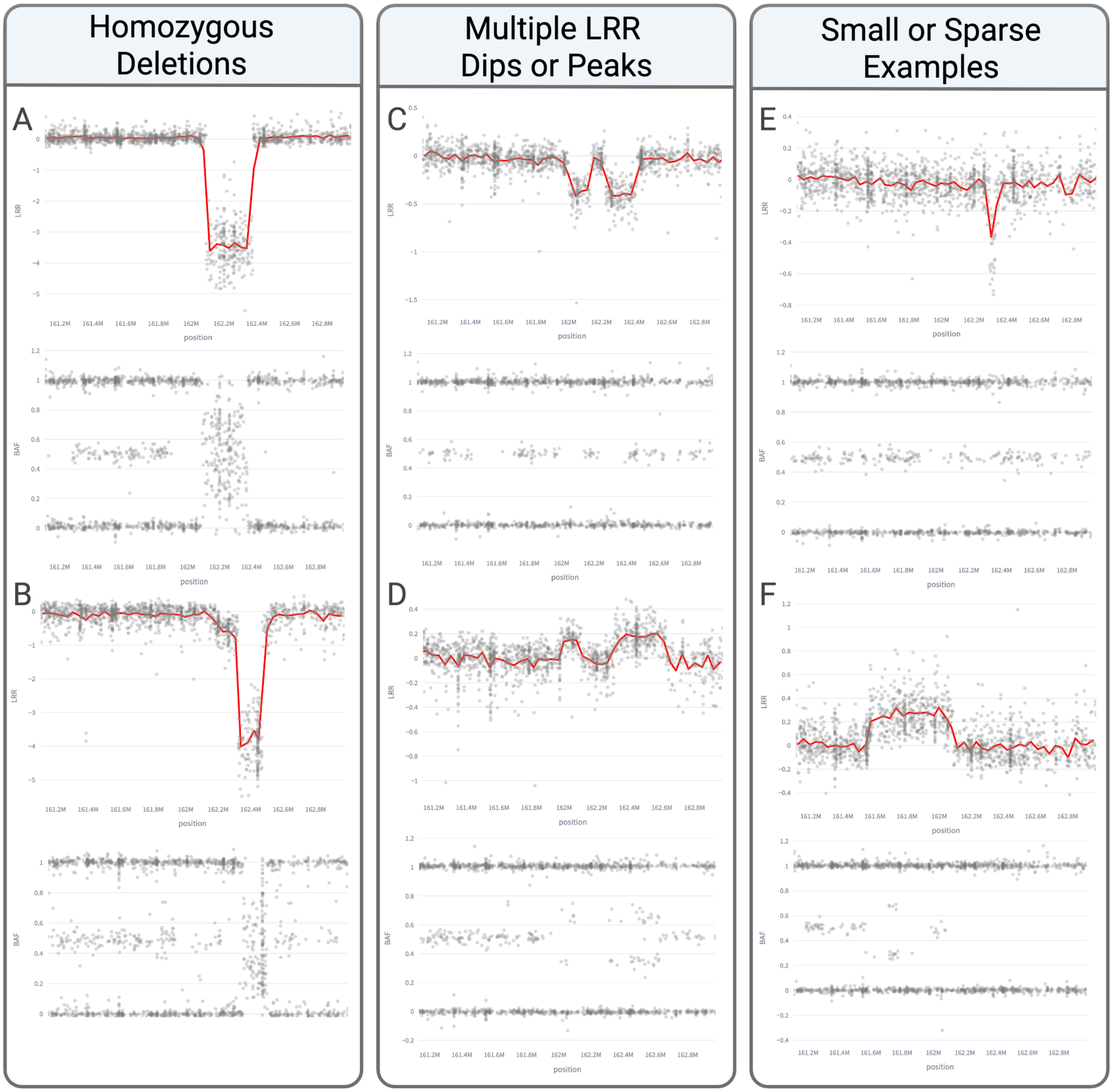


**Supplementary Figure 3**. Atypical samples identified by models without explicit training of these categories.

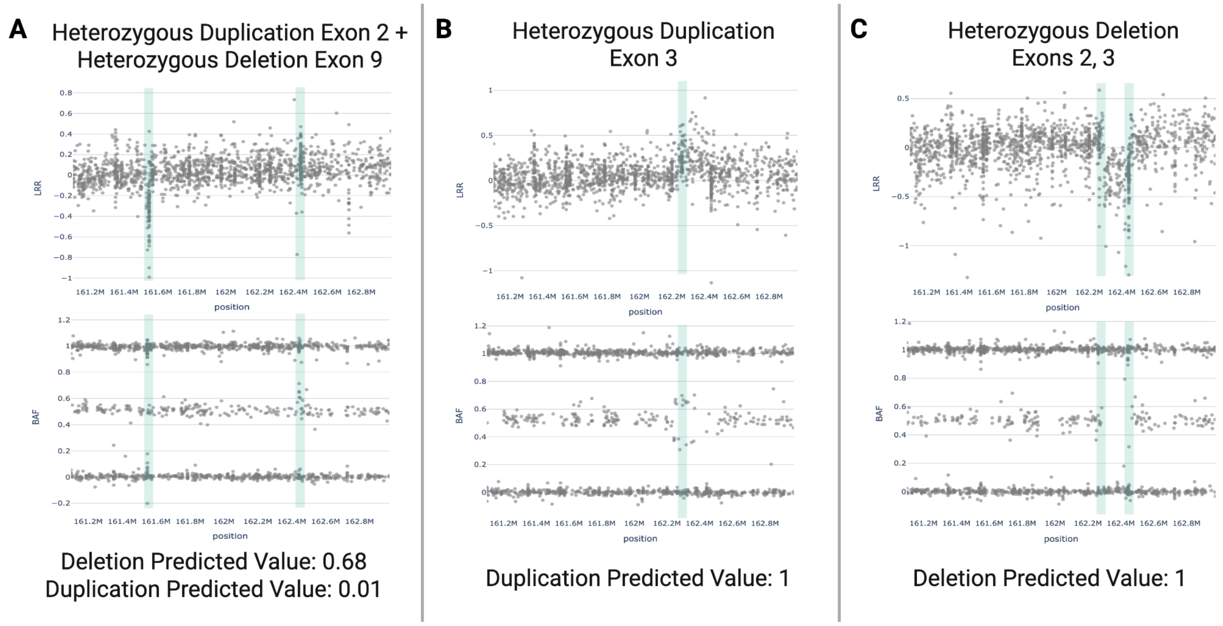


**Supplementary Figure 4**. Samples with MLPA hits in *PRKN* and the comparison of NBA-based plot features to the models' predicted values. Plots A and C feature Exon 2 which ranges from around 162,443,310-162,443,473 bp, Exon 9 in plot A spans approximately 161,548,854-161,549,003 bp, and Exon 3 in samples B and C includes positions 162,262,525-162,262,765 bp.

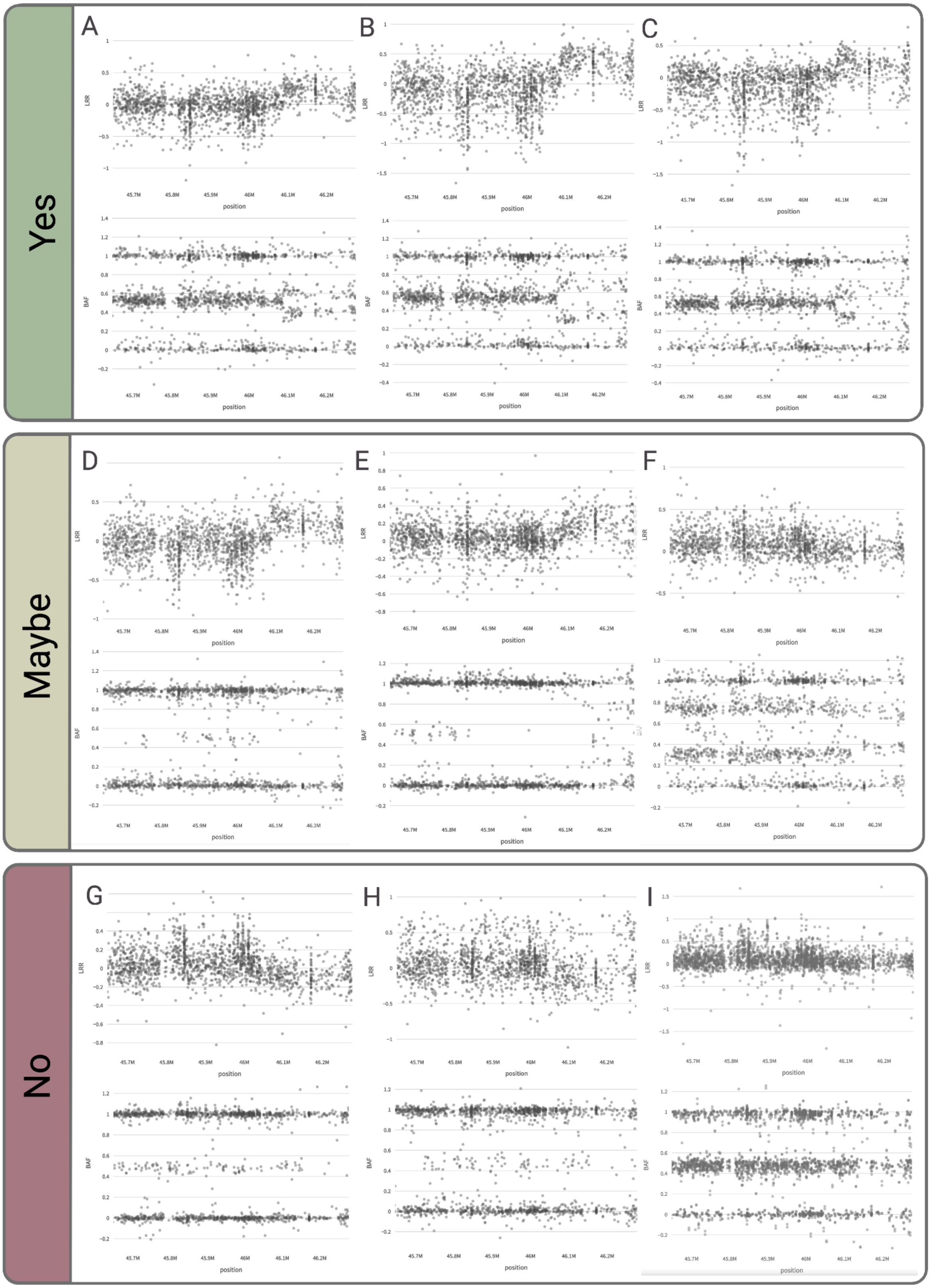


**Supplementary Figure 5**. Examples of classification used on duplication near *MAPT* during visual confirmation.
